# Genetic elements promote retention of extrachromosomal DNA in cancer cells

**DOI:** 10.1101/2025.10.10.681495

**Authors:** Venkat Sankar, King L. Hung, Aditi Gnanasekar, Ivy Tsz-Lo Wong, Quanming Shi, Katerina Kraft, Matthew G. Jones, Britney Jiayu He, Xiaowei Yan, Julia A. Belk, Kevin J. Liu, Sangya Agarwal, Sean K. Wang, Anton G. Henssen, Paul S. Mischel, Howard Y. Chang

## Abstract

Extrachromosomal DNA (ecDNA) is a prevalent and devastating form of oncogene amplification in cancer^1,2^. Circular megabase-sized ecDNAs lack centromeres and segregate stochastically during cell division^3–6^ yet persist over many generations. EcDNAs were first observed to hitchhike on mitotic chromosomes into daughter cell nuclei over 40 years ago with unknown mechanism^3,7^. Here we identify a family of human genomic elements, termed retention elements, that tether episomes to mitotic chromosomes to increase ecDNA transmission to daughter cells. We develop Retain-seq, a genome-scale assay that reveals thousands of human retention elements conferring generational persistence to heterologous episomes. Retention elements comprise a select set of CpG-rich gene promoters and act additively. Live-cell imaging and chromatin conformation capture show that retention elements physically interact with mitotic chromosomes at regions which are mitotically bookmarked by transcription factors and chromatin proteins, intermolecularly recapitulating promoter-enhancer interactions. Multiple retention elements are co-amplified with oncogenes on individual ecDNAs in human cancers and shape their sizes and structures. CpG-rich retention elements are focally hypomethylated; targeted cytosine methylation abrogates retention activity and leads to ecDNA loss, suggesting that methylation-sensitive interactions modulate episomal DNA retention. These results highlight the DNA elements and regulatory logic of mitotic ecDNA retention. Amplifications of retention elements promote the maintenance of oncogenic ecDNA across generations of cancer cells, revealing the principles of episome immortality intrinsic to the human genome.

## INTRODUCTION

Human cancer cells commonly amplify potent oncogenes on megabase-sized circular ecDNA molecules^8,9^ that lack centromeres and segregate asymmetrically^3–6^. This characteristic of ecDNA results in intra-clonal heterogeneity in oncogene copy number and amplicon sequence, as well as rapid adaptation to selective pressures during cancer evolution^6,8,10–12^. During cell division, the nuclear envelope breaks down before the segregation of chromosomes, which physically attach to the mitotic spindle at centromeres and partition into daughter nuclei. Thus, the acentric nature of ecDNA raises a crucial question: how is ecDNA inherited by daughter cells and retained within daughter nuclei after cell division? It has been well-documented that viral episomes such as papillomaviruses, Epstein-Barr virus (EBV), and simian virus 40 tether to mitotic chromosomes in order to hitchhike into daughter nuclei^13–17^. Viral episome tethering is mediated by dedicated viral DNA elements, viral DNA-binding proteins, and interactions with host cell chromatin-binding proteins, such as BRD4^13,18,19^. Notably, ecDNA strongly colocalizes with chromosomes during mitosis^3,20–23^, suggesting that ecDNA may also tether to chromosomes during DNA segregation. However, it is unknown what endogenous human DNA elements or factors mediate this tethering process. We hypothesized that such DNA sequences on ecDNA would enable it to be retained in the nuclear space of dividing cancer cells, thus serving as functional “retention elements”.

In principle, any ecDNA molecule that becomes amplified and persists in a cancer cell population should contain a minimum of three genetic elements: (1) a fitness element that provides a fitness advantage to the cell when under selective pressure (for example, an oncogene or regulatory sequence), (2) origins of replication to copy itself; and (3) a retention element that promotes nuclear retention of ecDNA by mediating its segregation along with chromosomes into daughter cells during cell division. In an evolving cancer cell population, ecDNA molecules with these features would become more abundant than molecules that lack them. While oncogenes^8,9,24^ and regulatory sequences^23,25,26^ on ecDNA as well as human origins of replication^27^ have been well studied, we have no understanding of the identity or mechanism of retention elements on human ecDNAs. Here, we devise a new genome-scale functional assay and apply imaging and chromatin profiling methods to elucidate the principles of genetic elements on ecDNA that promote its retention in dividing cells.

## RESULTS

### Genetic elements drive episome retention

We hypothesized that ecDNA is retained by hitchhiking onto chromosomes during cell division via the docking of ecDNA sites, which we term retention elements, to anchor sites on chromosomes (**Figure 1a**). We consider untethered ecDNAs (**Figure 1b**) as lost in this context, since acentric DNA that fails to segregate with chromosomes is released into the cytoplasm or incorporated into micronuclei^28–30^. This DNA is subject to strong transcriptional silencing, usually not replicated or expressed, and can be degraded and lost from the cell^30–32^. Live-cell time-lapse imaging of the colorectal cancer cells COLO320DM with ecDNA encoding the *MYC* oncogene (*ecMYC*) indeed showed synchronous segregation of ecDNA and chromosomal DNA during cell division (**Figure 1c**). Analysis of images of DNA fluorescence in-situ hybridization (FISH) paired with immunofluorescence (IF; IF-DNA-FISH) staining of Aurora Kinase B showed 97-98% colocalization of ecDNA with chromosomal DNA during segregation in multiple ecDNA-bearing cell lines (**Figure 1d**). These observations are consistent with past reports that ecDNA segregates synchronously with chromosomes and may tether to them^3,20–23^. Since these ecDNAs are derived from multiple distinct chromosomes, our results imply that functional retention elements must be widely dispersed across the human genome.

**Figure 1.**
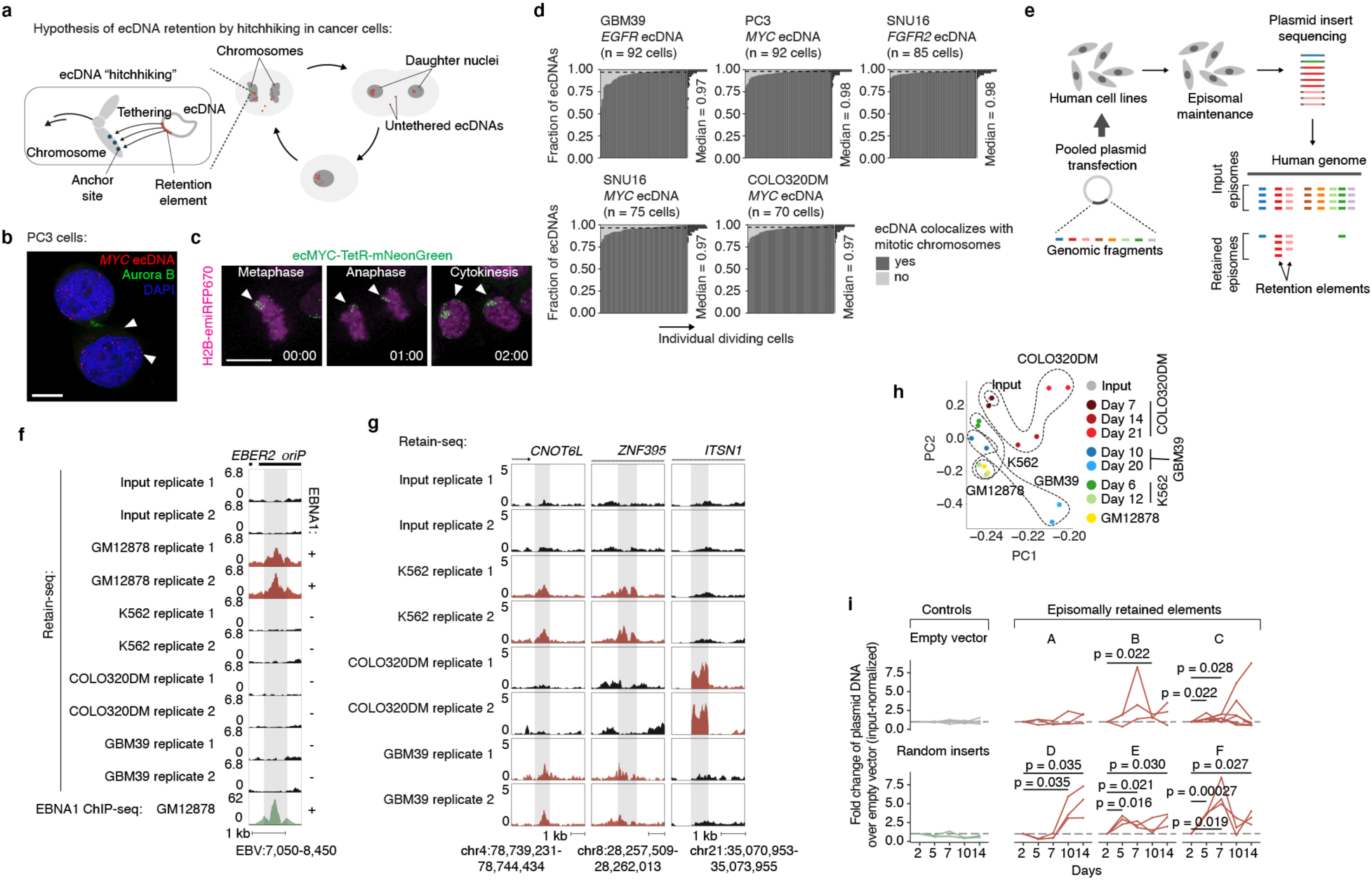
Identification of genetic elements that promote episomal DNA retention. **(a)** Hypothesis of mitotic retention of ecDNAs in cancer cells via chromosome hitchhiking. **(b)** Representative image of tethered (bottom arrow) and untethered (top arrow) ecDNA foci in mitotic PC3 cells (*n* = 92 daughter cell pairs). Scale bar, 10 µm. **(c)** Representative live-cell images (*n* = 10 fields of view) showing ecDNA (labeled with TetR-mNeonGreen) colocalization with chromosomes during cancer cell division. Scale bar, 10 µm. **(d)** Fractions of ecDNAs with various oncogenes colocalizing with mitotic chromosomes in cancer cell lines (glioblastoma GBM39, *EGFR* ecDNA from chromosome 7; prostate cancer PC3, *MYC* ecDNA from chromosome 8; gastric cancer SNU16, *MYC* and *FGFR2* ecDNAs from chromosome 8 and chromosome 10, respectively; colorectal cancer COLO320DM, *MYC* ecDNA (*ecMYC*); raw images obtained from a previous publication^5^) in IF-DNA-FISH of anaphase cells. **(e)** Schematic diagram of Retain-seq. **(f)** Retain-seq enrichment of a known EBV sequence that promotes viral retention, with EBNA-1 ChIP-seq in the EBV-transformed GM12878 cells below. **(g)** Retain-seq signal at three representative enriched genomic loci. Red tracks represent loci that were significantly enriched in Retain-seq screens in the corresponding cell line, thus marking these loci as retention elements in that line; black tracks indicate that the sequence was not identified as a retention element in the corresponding experiment. **(h)** Principal component analysis of Retain-seq in various cell lines at different time points. **(i)** Individual validation by quantitative PCR of six episomally retained elements identified by Retain-seq experiments in the K562 cell line and amplified on the COLO320DM (RE-C) and GBM39 (others) ecDNAs. Each line in the plot for a given retention element represents a single replicate. The empty vector control is the pUC19 plasmid alone, while the random inserts control comprises the pUC19 plasmid with random insert sequences from the genome of the human GM12878 cell line. *P*-values determined by one-sided t-test.

To broadly identify genetic sequences that may serve as retention elements on ecDNA, we developed a shotgun genetic screen, termed Retain-seq, that identifies episomally retained sequences (**Figure 1e**). Briefly, we created a pool of heterologous bacterial plasmids with inserts representing random DNA sequences from the human genome (**Figure 1e**, **Extended Data Figure 1a,b**). We then transfected the plasmid pool into multiple cell types, followed by serial passaging, and extracted retained plasmid DNA from the cells to identify enriched episomal DNA sequences by targeted sequencing of the inserts (**Figure 1e**). To minimize the effects of the variability of insert size and amount on the enrichment analysis due to PCR over-cycling, we halted PCR amplification at the cycle before saturation and performed all subsequent enrichment analyses comparing the output DNA with the transfected input episomal DNA library (**Extended Data Figure 1c,d**). A serial dilution experiment showed minimal over-representation of DNA sequences with variable amounts of DNA using this PCR strategy (**Extended Data Figure 1c**). As a validation for Retain-seq, we observed specific episomal enrichment of the oriP family of repeats (EBV:7,421-8,042), the EBV genomic sequence that enables viral tethering to chromosomes mediated by the virally encoded protein EBNA1^33^, only in the EBNA1-positive GM12878 cells but not in EBNA1-negative K562, COLO320DM, or GBM39 cells (**Figure 1f**). Retain-seq enrichment signal coincides strongly with EBNA1 occupancy (**Figure 1f**), consistent with the idea that EBNA1 binding to this viral element mediates episomal retention and tethering to chromosomes.

Next, we analyzed retained episomal DNA from multiple time points across two ecDNA-positive cell lines, COLO320DM and GBM39, and one ecDNA-negative cell line, K562 (**Figure 1g**). The sequence representation of the transfected library was comparable to that of the input episomal library; thus, the latter was used in subsequent analyses for identifying enriched elements (**Extended Data Figure 2a**). We then filtered out time points at which genome representation of the episomes dropped below our data quality threshold using the serial dilution experiment (**Extended Data Figure 2b**; Methods). Due to variation in transfection efficiency and growth rate across cell lines, we observed varying levels of stochastic drift in the retained episomal library between replicates over time (**Figure 1h, Extended Data Figure 2c**). To first capture retention elements with potential activity in any cell line, we identified a combined set of 14,353 retention elements (**Extended Data Figure 2d,e**). Most retention elements are captured within 1-kilobase (kb) genomic segments (**Extended Data Figure 2f**). To validate the ability of retention elements to retain episomal DNA in cells, we individually cloned six retention elements originally identified in the Retain-seq experiment in K562 cells into the pUC19 plasmid backbone and transfected these plasmids individually into K562 cells. These particular retention elements were chosen for validation because they also overlap with the coordinates of ecDNAs found in COLO320DM or GBM39 cells. Five out of six plasmids with retention elements were retained in K562 cells at higher levels compared to both the empty vector control and plasmids with random sequence inserts, validating the activity of retention elements identified by Retain-seq (**Figure 1i**). Although a subset of retention elements was both enriched and individually validated in multiple cell types (e.g., RE-C; **Figure 1i, 2j**), the majority appear unique to each cell type, reflecting cell type specificity or technical variation across cell lines. A positive control plasmid bearing the EBV tethering sequence alone displayed an increase in plasmid persistence of comparable magnitude relative to an empty vector control (**Extended Data Figure 2g**), showing that retention elements identified within the human genome promote episomal DNA retention to similar extents to known viral sequences. A retention element does not increase genomic integration of plasmids (**Extended Data Figure 3**), ruling out preferential integration of episomal elements into chromosomes as a mechanism of retention. Together, these results suggest that episomal retention elements are distributed broadly across the human genome.

### Retention elements comprise active DNA

We next sought to characterize the sequence features of retention elements (**Figure 2a**). We found that retention elements are highly enriched at transcription start sites (TSSs) and 5’ untranslated regions (UTRs) of genes (**Figure 2b,c**). By contrast, retention elements are depleted across the large stretches of distal intergenic regions (**Figure 2c**). Retention elements are broadly associated with regions of active chromatin, showing strong enrichment at gene promoters and enhancers (**Figure 2c,d**) and sites occupied by both actively elongating and paused RNA polymerase II (**Figure 2e**). As expected due to their overlap with promoter sequences, a substantial proportion of retention elements represent sites of nascent transcription (**Extended Data Figure 4a,b**). However, the presence of retention elements that are not actively transcribed and the fact that the majority of ecDNAs are maintained in the nucleus even after transcription inhibition by triptolide treatment^34^ suggest that transcription may not be necessary for function of all retention elements (**Extended Data Figure 4a,b**). Retention elements are also preferentially bound by the SWI/SNF chromatin remodeling complex, BRD4, CTCF, and histones with active marks such as H3K27ac, H3K4me3, and H3K9ac (**Figure 2e**, **Extended Data Figure 5a**). By contrast, retention elements show absence of overlap with RNA polymerase III or repressive histone marks such as H3K9me3 and H3K27me3 (**Figure 2e**). We found that CpG density is elevated in retention elements (**Figure 2f,g**), consistent with the idea that regions of active chromatin in the genome typically contain CpG-dense DNA sequences^35^. Because retention elements are CpG rich and do not appear heterochromatinized, they likely represent a separate class of sequences from AT-rich scaffold matrix attachment regions^36^ and rely on divergent protein factors for function. Importantly, we observed only minor overlap (∼8%) of retention elements with origins of replication and low occupancy of replication licensing complexes (MCM2-7) at retention elements, suggesting that retention elements do not promote episomal DNA enrichment by serving as origins of replication (**Figure 2h**, **Extended Data Figure 5b**). Furthermore, transfection of plasmids carrying either validated retention elements or a known EBV tethering sequence showed similar levels of retention in cells while inclusion of the full EBV origin, including a replicator sequence, dramatically increased plasmid DNA content by two orders of magnitude, supporting the conclusion that retention elements alone do not broadly induce DNA replication (**Extended Data Figure 2g**).

**Figure 2.**
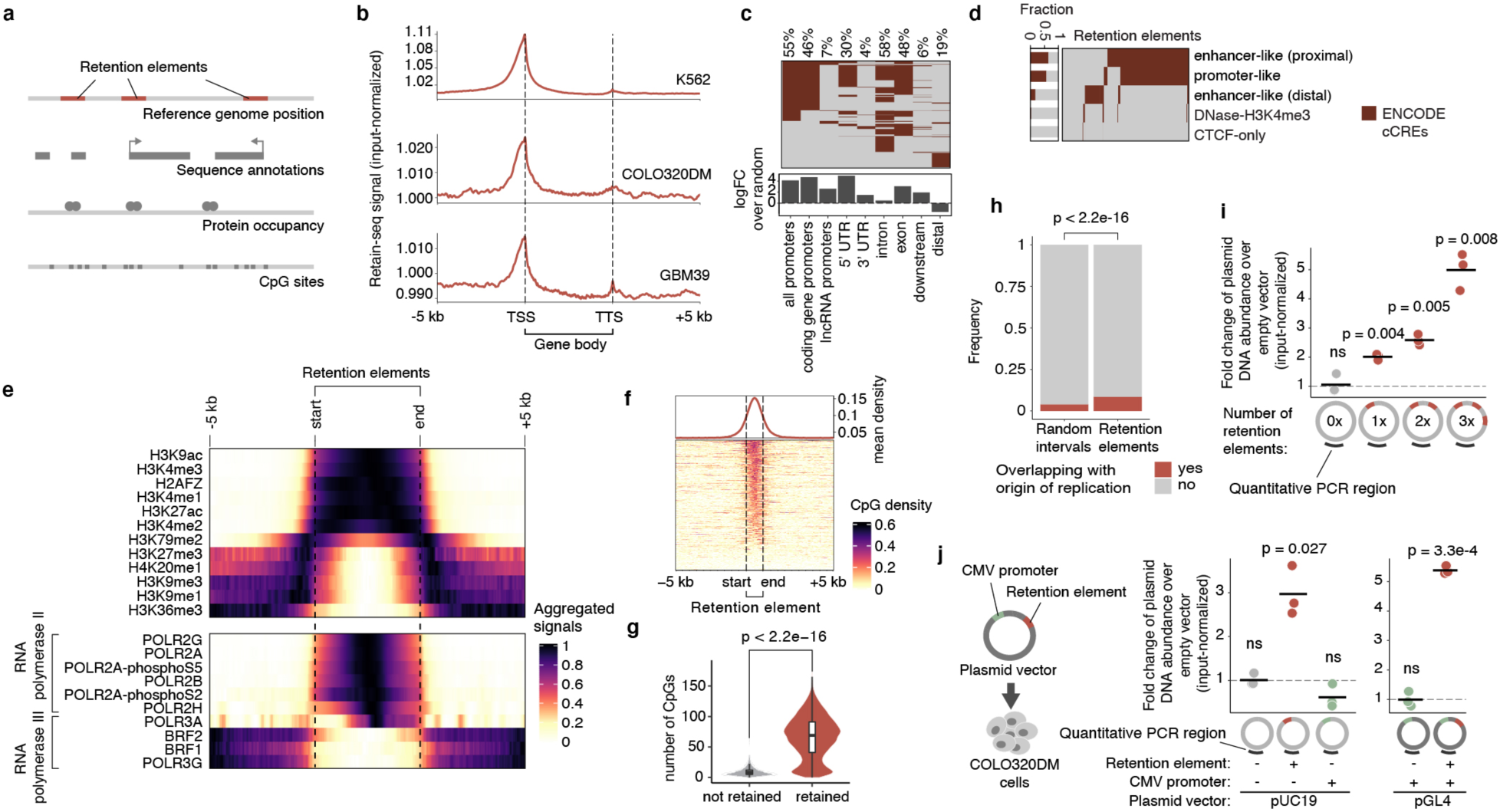
Sequence features of retention elements. **(a)** Analyses of sequence features of retention elements. **(b)** Input-normalized Retain-seq signal across annotated gene sequences. TSS, transcription start site; TTS, transcription termination site. **(c)** Sequence annotations overlapping with retention elements identified in K562 cells. Percentages represent the proportion of retention elements overlapping with a given annotation class. **(d)** ENCODE candidate cis-Regulatory Elements (cCREs) overlapping with retention elements identified in K562 cells. Fractions represent the proportion of retention elements overlapping with a given cCRE class. **(e)** ENCODE ChIP-seq signals of the indicated histone marks and RNA polymerase II and III in K562 cells surrounding retention elements identified in the same cell line. **(f)** CpG density surrounding the combined set of retention elements. **(g)** Number of CpG sites in genomic bins overlapping with retention elements (*n* = 18494) compared to those that do not (*n* = 2543727). Box center line median; limits, upper and lower quartiles; whiskers, 1.5× interquartile range. *P*-value computed by two-sided Wilcoxon rank-sums test. **(h)** Fraction of origins of replication (identified by SNS-seq in K562 cells) overlapping with retention elements identified in K562 cells and random genomic intervals. *P-*value determined by one-sided hypergeometric test. **(i)** Retention of plasmids containing one, two or three copies of a retention element (RE-C; red segments in schematic) in COLO320DM cells by quantitative PCR. Fold changes were computed using plasmid levels at day 14 post-transfection, normalizing to levels at day 2 to adjust for differential transfection efficiency across conditions (three biological replicates). *P-*values computed using one-sided t-test. **(j)** Left: transfection of plasmids with a CMV promoter and/or a retention element (RE-C) into COLO320DM cells. Right: retention of plasmids containing a CMV promoter and/or a retention element in COLO320DM cells by quantitative PCR (three biological replicates). Data for two different plasmid backbones, pUC19 and pGL4, are shown. *P-* values computed using one-sided t-test.

Episomal retention increased with the number of retention elements (**Figure 2i**). This additive effect also suggests that retention elements are functionally distinct from centromeres, as the presence of more than one centromere per episome or chromosome leads to opposing kinetochores pulling on the same DNA, leading to DNA fragmentation and loss^37^. Intriguingly, while we observed enrichment of gene promoters in retention elements (**Figure 2b-d**), the constitutive cytomegalovirus (CMV) promoter did not promote episomal retention alone (**Figure 2j**). This observation shows that an active promoter itself is not sufficient to enable DNA retention and suggests that additional sequence-specific interactions may be required. Consistent with this idea, we found that similar DNA motifs of chromatin-binding proteins are enriched across retention elements identified in multiple cell lines, suggesting that sequence features of retention elements may converge despite variation in the enriched intervals themselves across cell lines (**Extended Data Figure 5c**). As a preliminary effort to identify a minimal sequence sufficient for episomal retention, we split a retention element into 8 overlapping tiles and individually assayed each segment (**Extended Data Figure 5d**). However, no individual segment enabled episomal retention to the extent of the original larger sequence, suggesting a possible reliance on combinatorial interactions across multiple sites within this element (**Extended Data Figure 5d**). Together, these results show that retention elements are pervasive, additive, and functionally composite DNA elements.

### Retention elements tether to chromosomes

Next, we asked whether retention elements allow episomal DNA to tether to chromosomes during DNA segregation. Using the COLO320DM cell line with *ecMYC* edited to contain a Tet-operator (TetO) array, we introduced plasmid DNA containing a Lac-operator (LacO) array and assessed the localization of plasmid and ecDNA during DNA segregation using fluorescence labeling and live-cell imaging (**Figure 3a,b, Extended Data Figure 6a**). Plasmids bearing a retention element displayed significantly increased colocalization with chromosomes throughout mitosis compared to the empty vector control (**Figure 3c,d**). A single retention element more than halved the probability of failure of chromosome hitchhiking of the linked episome from 25% to 10.4% per mitotic event (**Figure 3c**). This difference was not observed in the TetO ecDNA signals between the two plasmid transfection conditions, validating uniform analysis across conditions (**Figure 3c,d**). This observation supports the idea that retention elements may increase episomal DNA retention by promoting its tethering to mitotic chromosomes. Ectopic plasmids with a retention element do not necessarily colocalize with endogenous ecDNAs (**Figure 3b, Extended Data Figure 6b,c**), indicating that retention elements confer autonomous retention activity.

**Figure 3.**
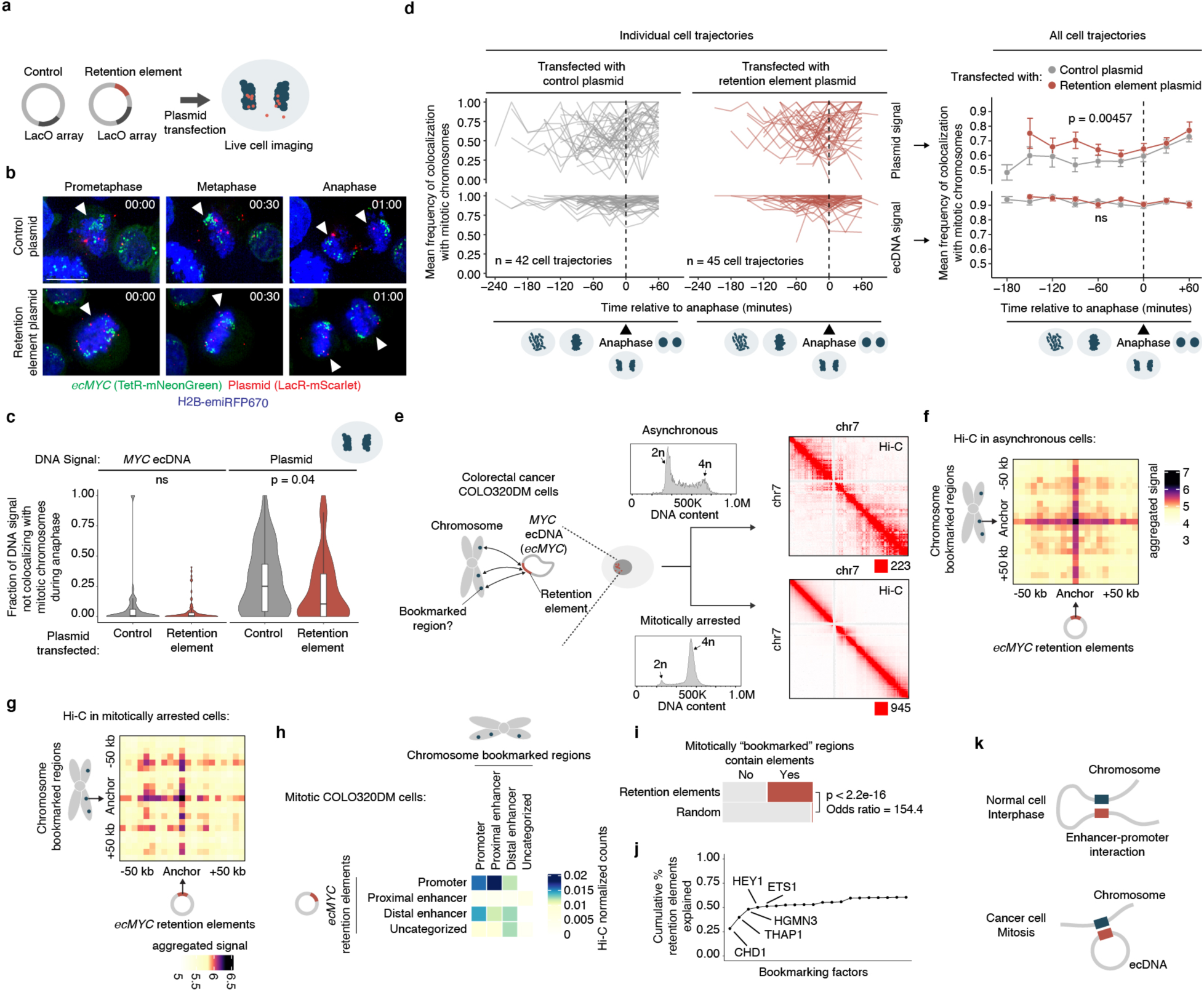
Retention elements promote extrachromosomal interactions with chromosomes during mitosis. **(a)** Live-cell imaging experiment schematic. **(b)** Representative live-cell time-lapse images of dividing COLO320DM cells with labeled *ecMYC* following transfection with plasmid containing a retention element or empty vector control. Scale bar, 10 µm. **(c)** Fraction of DNA signal not colocalizing with mitotic chromosomes during anaphase. *n* = 51 (control), *n* = 83 cells (retention element). Box plot parameters as in Fig. 2. *P*-values by two-sided Wilcoxon rank-sums test. **(d)** Individual (left) and mean (right) cell trajectories of DNA signal colocalization with chromosomes throughout mitosis. *n* = 42 (control), *n =* 45 (retention element) cells. Mean cell trajectories include all time points with > 3 cells. Error bars show s.e.m. *P*-values by two-sided paired t-test. **(e)** Hi-C interaction maps in asynchronous or mitotically arrested COLO320DM cells. Density plots show flow cytometric analysis of DNA content. **(f,g)** Aggregated peak analysis (APA) of Hi-C data of asynchronous (f) and mitotically arrested (g) COLO320DM cells. Heatmaps are summed percentile matrices of pairwise interactions between chromosome bookmarked regions and a combined set of *ecMYC* retention elements with 5-kb resolution. **(h)** Hi-C heatmap of pairwise interactions in mitotically arrested COLO320DM cells between *ecMYC* retention elements and chromosome bookmarked regions with ENCODE cCRE annotations. **(i)** Mitotically bookmarked regions overlapping with retention elements or matched-size random genomic intervals. *P*-values by two-sided Fisher’s Exact Test. **(j)** Cumulative distribution of retention elements containing binding sites of bookmarking factors, ordered by factor enrichment relative to random genomic intervals. **(k)** ecDNA-chromosome interactions recapitulate enhancer-promoter interactions. While gene expression in interphase cells is activated by an interaction between enhancer (blue) and promoter (red) sequences on the same chromosome, we hypothesize that ecDNA retention in mitotic cells is mediated by an analogous intermolecular contact between promoter-like retention elements (red) on ecDNA and enhancer-like, or less commonly, promoter-like bookmarked sites (blue) on the chromosome.

### Episomal contact with mitotic bookmarks

As our live-cell imaging analysis showed that a retention element promotes tethering of plasmids to chromosomes during mitosis, we asked whether retention elements on oncogene-carrying ecDNAs in cancer cells (i.e., genomic intervals within the ecDNA that coincide with retention element intervals identified by Retain-seq) may contact specific sites on chromosomes. While chromosomes are compacted 10,000-fold during mitosis, some genomic sites remain accessible and are stably bound by transcription factors throughout mitosis^38–44^, a phenomenon termed “mitotic bookmarking”. To first interrogate whether ecDNA-chromosome interactions occur at mitotically bookmarked loci, we performed genome-wide chromatin conformation capture using Hi-C on mitotically arrested COLO320DM cells to analyze pairwise DNA interactions between *ecMYC* and chromosomes (**Figure 3e**). As expected, pairwise chromatin interaction maps showed plaid patterns of long-range interactions in asynchronous cells but substantial loss of these long-range interactions in mitotically arrested cells due to chromatin condensation (**Figure 3e**), consistent with previous Hi-C studies^45^. Next, we performed aggregate peak analysis (APA) to measure enrichment of Hi-C signal in pairs of loci, with one partner on *ecMYC* containing a retention element and the other partner on a chromosome containing a mitotically bookmarked region (**Figure 3f,g**). We observed enrichment of Hi-C contacts between chromosome bookmarked regions and ec*MYC* retention elements in asynchronous cells, which are retained in the condensed chromatin of mitotically arrested cells despite increased background noise (**Figure 3f,g**). By contrast, we did not observe focal interactions when either or both the chromosomal or extrachromosomal regions were randomized (**Extended Data Figure 7a,b**). These data suggest that focal interactions occur between retention elements on ecDNA and mitotically bookmarked regions on chromosomes both in interphase and during mitosis. This behavior is analogous to that of the EBV episomal genome, which also remains associated with chromosomes throughout the cell cycle^33^. The majority of chromosome bookmarked regions overlap with promoters or proximal enhancer-like elements, while *ecMYC* retention elements consist of distal enhancer-like elements and promoters (**Extended Data Figure 7c**). Notably, retention elements on *ecMYC* overlapping with promoters showed increased Hi-C contact with proximal enhancer-like elements and promoters at chromosome bookmarked regions, while retention elements on *ecMYC* overlapping with distal enhancer-like elements showed increased Hi-C contact with chromosome bookmarked loci originating from promoters (**Figure 3h, Extended Data Figure 7d**). We also performed APA on Hi-C data from asynchronous GBM39 cells, though this analysis was inconclusive likely due to a small sampling size as the ecDNA of this cell line contains a smaller number of retention elements (**Extended Data Figure 7e**).

Because factors promoting ecDNA retention via chromosomal hitchhiking should bind to condensed chromosomes during mitosis, mitotic bookmarking factors are plausible candidates as mediators of ecDNA retention. Nearly half of the mitotically bookmarked regions were also identified as retention elements, which is highly enriched over randomly selected genomic intervals of the same size (**Figure 3i**). Many putative bookmarking factors represented by ChIP-seq data in K562 cells (ENCODE consortium^46^) showed occupancy within retention elements, with as few as five bookmarking factors cumulatively binding over 50% of retention element intervals (**Figure 3j**). Intriguingly, a subset of bookmarking factors consistently bound more retention elements than others, indicating that some factors may disproportionately contribute to retention element activity (**Extended Data Figure 7f**). However, individual CRISPR-mediated knockouts of three enriched bookmarking factors did not result in widespread untethering of ecDNA in mitotic COLO320DM cells, suggesting that mitotic ecDNA retention involves complexes of multiple redundant DNA binding proteins on active chromatin^47^ (**Extended Data Figure 7g,h**). Together, these observations support the idea that ecDNA-chromosome interactions in mitotic cancer cells intermolecularly recapitulate promoter-enhancer interactions (**Figure 3k**).

### Cancer ecDNAs contain retention elements

While retention elements promote the maintenance of episomal DNA in dividing cells, ecDNAs also provide selective advantages to cancer cells by encoding oncogenes. Thus, ecDNAs can theoretically become amplified in a cell population due to selection despite imperfect retention during cell division. To explore the relative contributions of retention and selection on ecDNA amplification, we simulated growing cancer cell populations by adapting an evolutionary framework^6^ to model imperfect retention. While ecDNAs were amplified with increased selection as expected, they were rapidly lost when the retention fidelity of ecDNAs per cell division dropped below 0.9 (**Figure 4a, Extended Data Figure 8a**), suggesting that a very high level of mitotic retention is a pre-requisite for selection to drive ecDNA amplification. Intriguingly, this minimum predicted level matches the experimentally observed mitotic retention rate (10% failure rate per mitosis) conferred by a single retention element based on live cell imaging (**Figure 3c**). Mitotic retention remains crucial even after ecDNAs reach high copy numbers, as imperfect retention led to loss of ecDNAs over time even in cells that have already reached high copy numbers and in the presence of selection (**Extended Data Figure 8b**).

**Figure 4.**
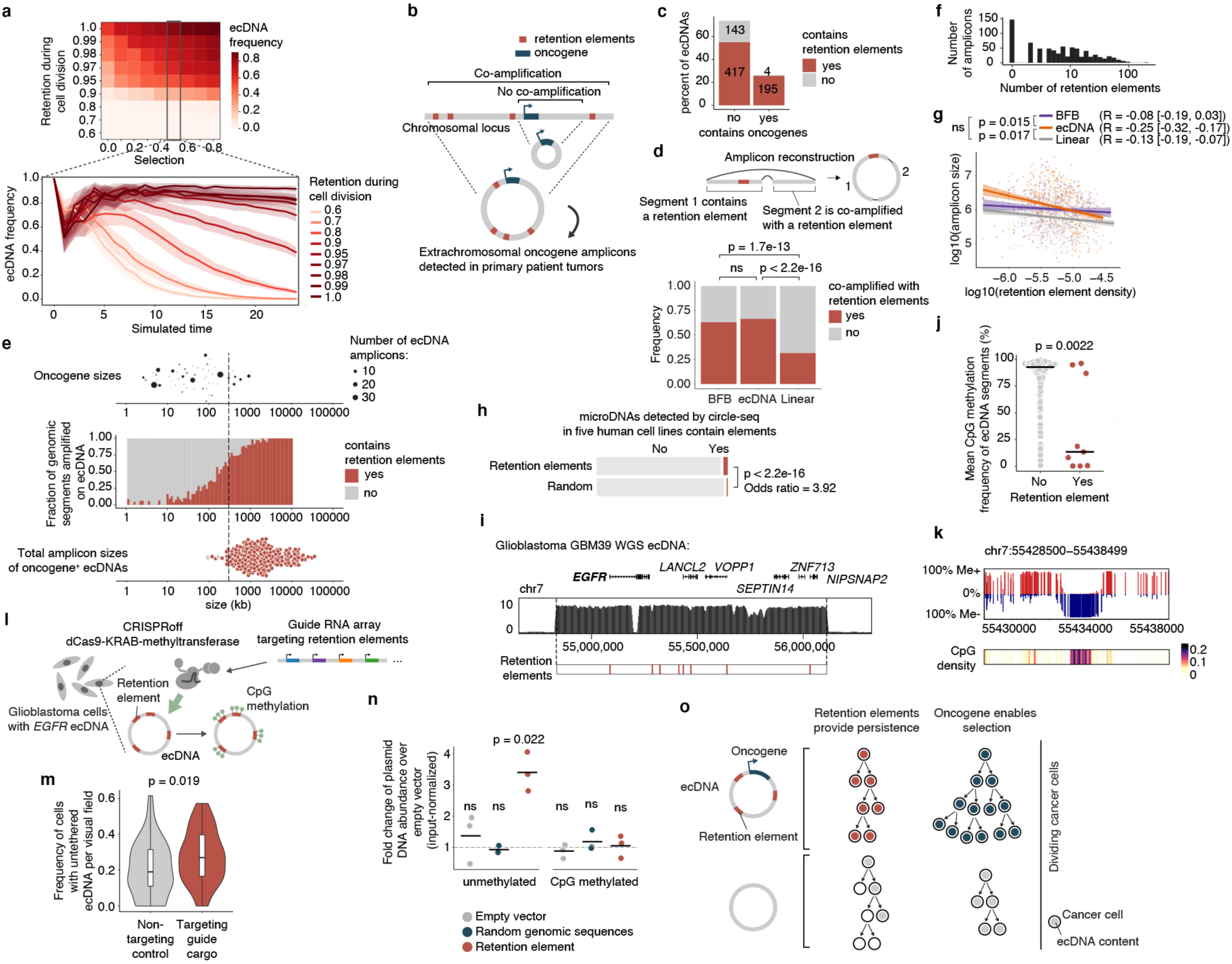
Retention elements enable selection of oncogene-carrying ecDNAs in cancer. **(a)** Mean frequency (over 10 independent replicates) of cells carrying ≥1 ecDNA in simulations. Shaded area, s.e.m. **(b)** Analysis of retention element co-amplification with oncogenes on ecDNA in patient tumors. **(c)** ecDNA amplicons containing retention elements and/or oncogenes. **(d)** Top: an ecDNA segment lacking retention elements co-amplified with a retention element. Bottom: frequency of co-amplification with retention elements within BFB, ecDNA, or linear amplicons for genomic segments lacking retention elements. One-sided test of equal proportions. **(e)** Top to bottom: oncogene sizes on ecDNA; frequency of genomic segments containing retention elements sorted by size; total ecDNA amplicon sizes. **(f)** Distribution of retention element numbers among ecDNAs. **(g)** Correlation (Pearson’s *R*; 95% confidence intervals) between local density of retention elements (Methods) and amplicon size. *P*-values by two-sided Fisher’s z-test. Plot: Linear fit (OLS) with 95% confidence intervals. **(h)** Circular microDNAs in five human cell lines overlapping with retention elements or matched-size random genomic intervals. Two-sided Fisher’s Exact Test. **(i)** Elevated WGS coverage of *EGFR* ecDNA in GBM39 cells and retention element positions. **(j)** 5mC CpG methylation of retention elements (*n* = 9 segments) compared to matched-size sequence intervals (*n* = 1235 segments) within the GBM39 ecDNA. Two-sided Wilcoxon rank-sums test. **(k)** 5mC methylation and density of CpG sites surrounding a retention element on the GBM39 ecDNA. **(l)** Site-specific methylation of retention elements by CRISPRoff. **(m)** Frequency of GBM39 cells containing untethered ecDNA foci 5 days after transfection*. n* = 60 (non-targeting) and *n* = 50 (targeting) visual fields. Box plot parameters as in Fig. 2. Two-sided Mann-Whitney-Wilcoxon test. **(n)** Plasmid retention after methylation in COLO320DM cells by quantitative PCR (three biological replicates). One-sided t-test. **(o)** Retention elements and oncogenes on ecDNA (left) confer retention and selection, respectively, two processes shaping the evolution of cancer cell lineages (right).

We next asked whether copy-number amplified, oncogene-carrying ecDNAs from patient tumor samples contain retention elements (**Figure 4b**). Analysis of focal amplifications in whole-genome sequencing (WGS) data from two patient cohorts (**Extended Data Figure 9a**) revealed that nearly all oncogene-containing ecDNAs contain retention elements (98%, **Figure 4c**). DNA segments that do not contain retention elements are often connected with those containing retention elements on ecDNAs but not chromosomal linear amplicons, even after adjusting for rearrangement events (**Figure 4d, Extended Data Figure 9b**). Breakage fusion bridge (BFB) amplifications, which can generate both ecDNAs and complex linear amplicons, also show similar enrichment of retention element co-amplification (**Figure 4d**). Moreover, observed ecDNAs are ∼10-fold larger in size (>1 megabase) than the oncogene-coding sequences and their cognate regulatory elements (∼100 kb); thus, nearly all observed ecDNA sequence coordinates encompass large segments of additional DNA sequence to reach megabase-scale sizes at which they are very likely to contain multiple retention elements (**Figure 4e,f**), which serially increase the likelihood of extrachromosomal maintenance (**Figure 2i**). By contrast, linear amplicons cover a more dispersed range of sizes, frequently containing smaller amplicons that are less likely to contain retention elements (**Extended Data Figure 9c-d**).

To address whether the distribution of retention elements near an oncogene shapes the amplification of DNA sequence, we analyzed the degree of co-amplification between each specific retention element and each of two oncogenes frequently amplified on ecDNA, *EGFR* and *CDK4* (**Extended Data Figure 9e**). We observed skewing of ecDNA amplicon distributions in the non-coding regions containing retention elements upstream of the oncogene promoters (**Extended Data Figure 9f**). Selection for large amplicons may be explained by either inclusion of retention elements or co-amplification of distal enhancers^25^. However, examining the distributions of retention elements across all ecDNA loci, we found that amplicon size decreases as the local density of retention elements increases (**Figure 4g**), suggesting that retention-element-sparse regions of the genome are selected with larger ecDNA sequences that are more likely to capture retention elements, whereas smaller ecDNA sequences are selected in retention-element-dense regions. This relationship is observed to a significantly greater extent in ecDNAs compared to linear amplicons (**Figure 4g**) across a broad diversity of cancer types expressing varied oncogenes. These results support the premise that co-amplification of multiple retention elements with oncogenes on ecDNAs provides a selective advantage and shapes ecDNA structure.

While large clonally selected ecDNAs are frequently observed in cancer, small (sub-kilobase-sized) non-clonal extrachromosomal circular DNAs (eccDNAs, also termed microDNAs) that often lack gene-coding sequences have been detected in healthy somatic tissues^48,49^. These microDNAs are not maintained at amplified copy numbers and result from DNA fragmentation from across the entire genome^48^. The vast majority (96.5%) of microDNAs lack retention elements, as expected; nonetheless, we observed an enrichment for retention elements in observed microDNA sequences (LNCaP, C4-2, PC-3, OVCAR8 and ES-2 cell lines; previously published^50^), consistent with the idea that extrachromosomal DNA which contains retention elements may be more persistent in cells (**Figure 4h**). Collectively, these results show that the distribution of retention elements in the genome shapes the presence and sequence of DNA outside chromosomes.

### Methylation silences retention elements

As retention elements are CpG-rich promoters and associate with chromosomal bookmarked regulatory elements, we hypothesized that cytosine methylation of these CpG sites, known to silence promoter activity and inhibit transcription factor binding^51^, may affect interactions between retention elements and cellular components that promote their retention. We found that retention elements on ecDNA are hypomethylated (**Figure 4i-k**). Six of nine candidate retention element intervals in the *EGFR* ecDNA in GBM39 glioblastoma neurospheres are significantly demethylated compared to all other sequence intervals of 1 kb width on the same ecDNA (**Figure 4j**). Analysis of the *EGFR* ecDNA in GBM39 cells by single-molecule long-read sequencing^12^ confirmed specific and focal hypomethylation at retention elements (**Figure 4j,k, Extended Data Figure 10a**). To test whether CpG methylation impacts ecDNA retention, we used a catalytically-dead Cas9 fused to DNA methyltransferase (CRISPRoff^52^) to program site-specific CpG methylation simultaneously on five hypomethylated retention elements on the *EGFR* ecDNA in GBM39 neurospheres (**Figure 4l**; Methods). Targeted methylation of retention elements dramatically reduced growth and viability of GBM39 cells, as expected following loss and silencing of the ecDNA-encoded oncogenes that are key drivers of cancer cell survival (**Extended Data Figure 10b,c**). Due to the acute loss of viability in cells with ecDNA retention elements targeted by CRISPRoff, we were limited to collecting cells at early time points and did not observe a reduction in total ecDNA copy number at 5 days post-transfection (**Extended Data Figure 10d**). However, turning to imaging to isolate ecDNA tethering from the effects of oncogene silencing, we found that CRISPRoff targeting of retention elements significantly increased the frequency of cells with untethered ecDNA foci and reduced nuclear ecDNA compared to non-targeting controls (**Figure 4m, Extended Data Figure 10e,f**). To further ensure that ecDNA depletion is due to silencing of retention element function rather than negative selection due to transcriptional silencing of the oncogene, we leveraged our episome retention assay. *In vitro* CpG methylation of a plasmid containing a single retention element, but no coding genes, completely ablates the episomal retention conferred by this genetic element (**Figure 4n**). We corroborated these data by live cell imaging, independently showing that methylation decreased physical colocalization of plasmid DNA with mitotic chromosomes during DNA segregation (**Extended Data Figure 10g**). Together, our results show that episomal retention of DNA is promoted by retention elements whose hypomethylation at CpG sites not only augments oncogene transcription, but also enables the molecular interactions required to confer retention of episomal DNA.

## DISCUSSION

EcDNAs are powerful drivers of oncogene expression in human cancers but live with the mortal risk of being lost with every cell division. Ensuring its faithful transmission into daughter cells is an evolutionary imperative to achieve “episome immortality”. Through genome-wide functional screening, imaging and chromatin profiling, we discovered a new class of pervasive genomic elements that promote retention of extrachromosomal DNA copies in dividing cells (**Figure 4o**). We have shown that these retention elements comprise transcriptionally active regions of the human genome and are co-amplified on oncogenic ecDNAs in human cancers. Retention elements physically interact with mitotically bookmarked regions on chromosomes and promote tethering of extrachromosomal DNA to chromosomes during mitosis. Furthermore, the extrachromosomal retention of these genomic elements is sensitive to methylation at CpG sites, suggesting that molecular interactions that mediate DNA retention can be perturbed via epigenetic modifications. As ecDNA molecules that contain retention elements should in theory outcompete those that lack them in a cancer cell population, ecDNA retention likely represents a selection process that shapes the size and sequence of amplified DNA in cancer genomes.

We introduce Retain-seq as a mechanism-agnostic platform to discover functional DNA retention elements in human cells. We showed with live cell imaging that inclusion of a retention element can promote colocalization of episomal DNA with mitotic chromosomes. This result is consistent with the idea that tethering of acentric DNA to chromosomes promotes its retention in the nuclear space of dividing cells. However, we do not rule out orthogonal mechanisms^53^ by which ecDNA can be retained in cells. We recently reported the phenomenon of ecDNA co-segregation, in which multiple ecDNA species in a cell can be co-inherited by the same daughter cell during cell division^6^. Concomitant with intermolecular interactions between ecDNA species that facilitate their co-segregation, ecDNA hitchhiking may also occur indirectly if an ecDNA interacts with another ecDNA that contains retention elements. As the composition of retention elements encoded in the ecDNA amplicon may impact the fidelity of its inheritance, the sequence compositions and sizes of ecDNA species are likely a source of variation among ecDNA species and cancer cells.

Our results suggest that retention elements repurpose long-range DNA contacts via mitotic bookmarking for ecDNA hitchhiking. In interphase cells, interactions between enhancers and promoters allow multiple DNA regulatory elements to contact and activate genes up to 1 Mb away on the linear chromosome, typically in *cis* on the same chromosome. Large condensates that include Mediator and RNA polymerase II maintain this linkage, enabling active transcription^54,55^. During mitosis, transcription is silenced and transcription factors dissociate from condensed mitotic chromosomes. However, certain transcription factors and chromatin-binding proteins are retained, allowing prompt resumption of gene expression and cell fate in the daughter cells. Rather than a binary classification, recent studies indicate that many transcription factors continue to dynamically interact with mitotic chromosomes, and mitotic bookmarking factors enjoy longer occupancy time on mitotic chromosomes^38–44^. Thus, ecDNA may tether to chromosomes during mitosis by recapitulating long-range contacts between bookmarked enhancers and promoters, but in *trans* across distinct DNA molecules. The repurposing of mitotic bookmarks explains why retention elements are pervasive throughout the human genome, and suggests that many if not most chromosomal segments sufficiently large are capable of becoming persistent ecDNAs, provided that they confer selective advantages to cells. Intriguingly unlike chromosomes, ecDNAs possess highly accessible chromatin^56^ and continue to transcribe RNA at the onset of mitosis^6^, which may promote retention^47^. EBV and papillomavirus episomes bind BRD4^18,57^ and yeast selfish 2 micron plasmids bind the SWI/SNF complex^58^ to hitchhike on mitotic chromosomes; both BRD4 and SWI/SNF are prominent mitotic bookmarks^59,60^, suggesting a unifying principle. Our discovery that human retention elements require DNA demethylation suggests ecDNA selection occurs both at the genetic level for oncogene cargo and at the epigenetic level for active retention element states. We are inclined to believe that the more a retention element is active as a promoter and demethylated in its native chromosomal context, the more likely that such element can facilitate retention when liberated as ecDNA. Future systematic functional studies may identify factors that are necessary for ecDNA hitchhiking and interrogate the generalizability of retention element behavior across varied cell types. Identification of these mediators of ecDNA retention may enable the design of novel cancer therapies targeting the maintenance of oncogene copies.

Together, our work illustrates how a new class of genomic elements promotes the retention of ecDNA in actively dividing cancer cells. These genomic elements may drive selection of amplicon sequences and structures in cancer, impacting the process of DNA amplification and evolutionary trajectories of cancer clones. A mechanistic understanding of ecDNA retention may provide insights about how different cancer cell populations adopt various levels of oncogene copy number changes and how specific ecDNA amplicon sequences are selected in tumors. Beyond oncogene amplification in cancer, our model of extrachromosomal retention of DNA sequences may provide a general framework for understanding the minimal unit of DNA maintenance in human cells and guide the design of synthetic DNA cargos for cellular engineering efforts.

## Acknowledgements

We thank the members of the Chang and Mischel labs for discussion and C. Luong for assistance with Hi-C data mapping. This project was supported by Cancer Grand Challenges CGCSDF-2021\100007 with support from Cancer Research UK and the National Cancer Institute (H.Y.C., P.S.M.). V.S. was supported by an NSF Graduate Research Fellowship (DGE-2146755). K.L.H. was supported by a Stanford Graduate Fellowship and an NCI Predoctoral to Postdoctoral Fellow Transition Award (NIH F99CA274692 and K00CA274692). M.G.J. was supported by an NCI Pathway to Independence Award (NIH K99CA286968). X.Y. is a Damon Runyon Fellow supported by the Damon Runyon Cancer Research Foundation (DRG-2474-22). J.A.B. was supported by an HHMI Hanna Gray Fellowship. H.Y.C. was an Investigator of the Howard Hughes Medical Institute.

## Author Contributions

K.L.H. and H.Y.C. conceived the project. V.S., K.L.H., P.S.M., and H.Y.C. designed the study. V.S. and K.L.H. optimized the Retain-seq protocol, collected and analyzed Retain-seq data, performed quantitative PCR validations of retention elements via individual plasmid transfections, analyzed retention element sequence features, integrated ENCODE data, analyzed retention elements in patient tumor sequencing data, and performed experiments involving in vitro CpG methylation of plasmids. V.S. performed plasmid transfection experiments testing EBV elements, multiple copies of retention elements, sub-tiling of a retention element, and the effects of the CMV promoter in different plasmid vectors, and performed nanopore sequencing of cells transfected with plasmids to assess genomic integration. K.L.H. developed Retain-seq, analyzed IF-DNA-FISH, analyzed Hi-C data, integrated bookmarking factor analyses, and analyzed CpG methylation in nanopore sequencing data. V.S., K.L.H., Q.S., B.J.H., K.J.L. and S.K.W. performed molecular cloning of plasmids. A.G., I.T.L.W., V.S., X.Y., Q.S., and S.A. prepared reagents for and performed live-cell imaging. A.G., K.L.H., and I.T.L.W. processed and analyzed live-cell imaging data. K.K. performed mitotic arrest and prepared Hi-C libraries. V.S. performed CRISPR/Cas9 knockouts of bookmarking factors. I.T.L.W. and K.L.H. processed and analyzed mitotic IF-DNA-FISH imaging data from knockout cells. M.G.J. performed evolutionary modeling of ecDNA retention and selection. V.S., Q.S., J.A.B., and K.L.H. designed and performed CRISPRoff experiments. A.G., I.T.L.W., and V.S. processed and analyzed CRISPRoff imaging data. A.G.H., P.S.M., and H.Y.C. guided data analysis and provided feedback on experimental design. V.S., K.L.H., and H.Y.C. wrote the manuscript with input from all authors.

## EXTENDED DATA FIGURES

**Extended Data Figure 1.**
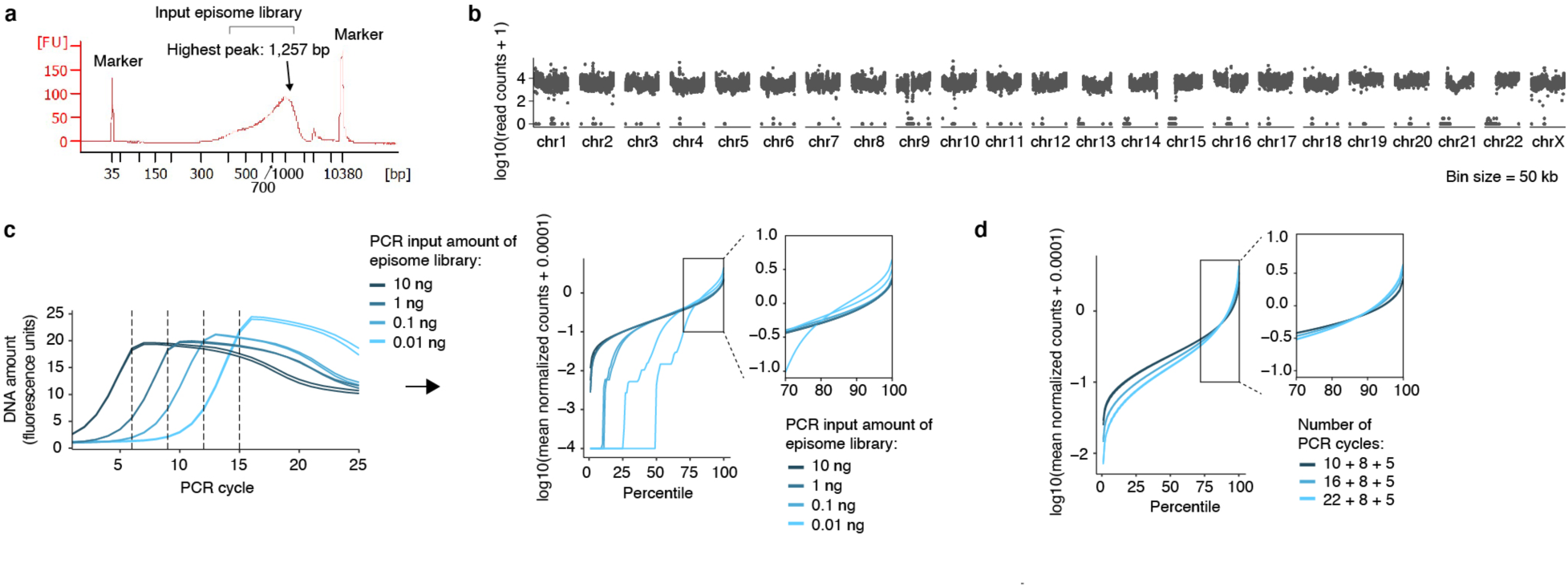
Optimization of Retain-seq library preparation. **(a)** Insert size distribution of genomic fragments included in the input mixed episome library. **(b)** Genome-wide coverage of sequenced reads derived from input episome library. **(c)** Left: Representative quantitative PCR amplification curves across varying amounts of episome library as PCR input. Right: Log-transformed mean normalized read counts of genomic bins ranked by percentile. Inset is a zoom-in of the higher-percentile genomic bins, in which a 100-fold range of DNA amounts from 0.1 ng – 10 ng of input showed highly comparable representation (despite some library dropout at 0.1 ng of input DNA) while 0.01 ng PCR input showed substantial library dropout and signs of skewing and was used to set the quality threshold for all library preparations. See Methods. **(d)** Log-transformed mean normalized read counts of genomic bins ranked by percentile. Inset is a zoom-in of the higher-percentile genomic bins showing that increasing PCR cycles during library preparation alters skewing of sequencing reads.

**Extended Data Figure 2.**
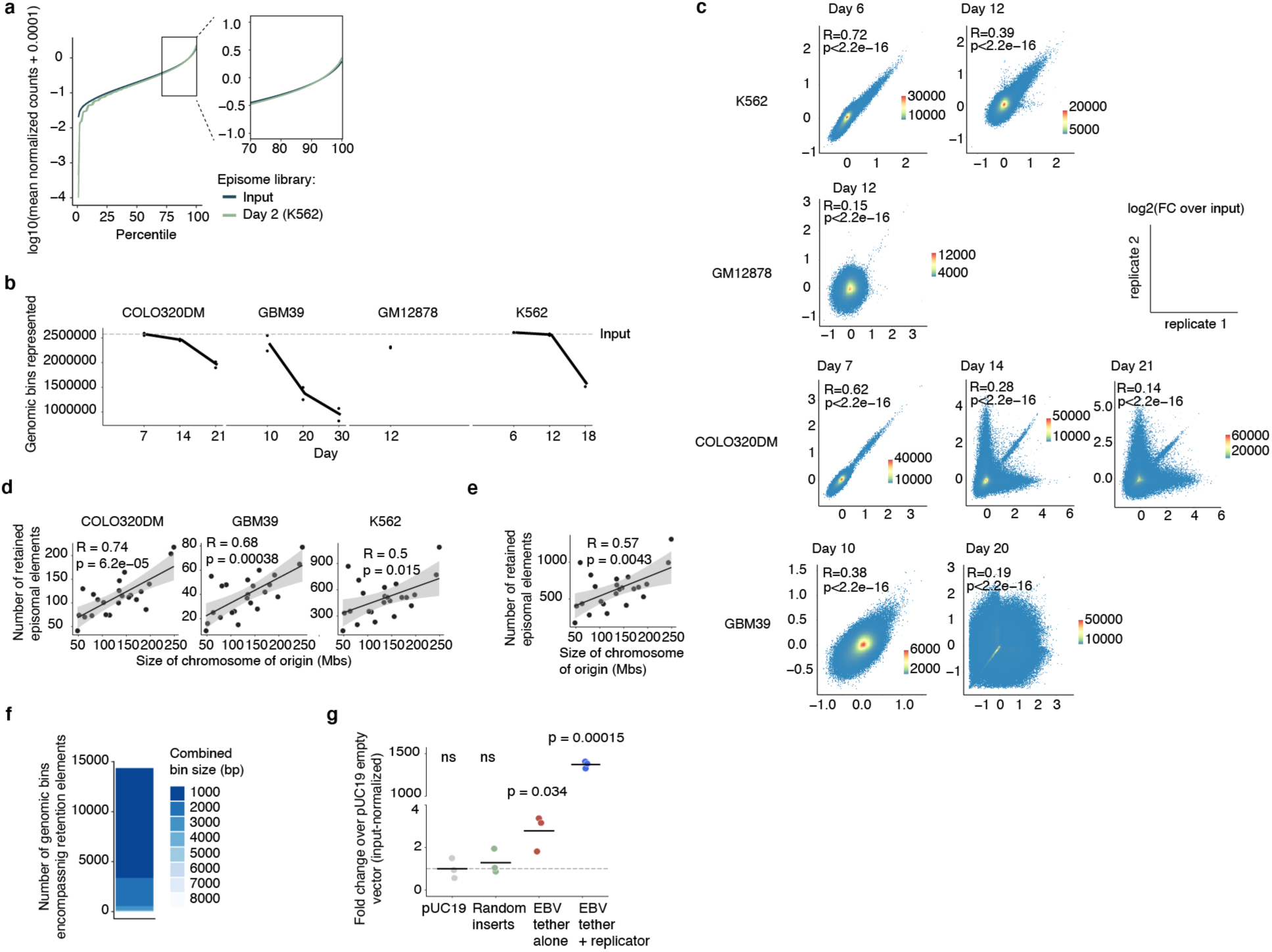
Distribution of Retain-seq reads across the genome and experimental replicates. **(a)** Log-transformed mean normalized read counts of genomic bins ranked by percentile. Inset is a zoom-in of higher-percentile genomic bins showing that transfection, represented by the day 2 episome library, results in minimal dropout that does not substantially skew the sequence representation compared to the input episomal library. **(b)** Loss of genome-wide representation in episomal insert sequences relative to the input library over time in four cell lines assayed with Retain-seq. **(c)** Correlations between experimental replicates of Retain-seq across time points from different cell lines. **(d)** Correlation (Pearson’s *R*; error bands represent 95% confidence intervals) between the numbers of episomally retained elements and the sizes of their chromosomes of origin in experiments performed in various cell lines. **(e)** Correlation (Pearson’s *R*; error bands represent 95% confidence intervals) between the numbers of episomally retained elements and the sizes of their chromosomes of origin across all cell lines. **(f)** Distribution of genomic bin sizes containing retention elements (median 1 kb; s.d. 0.604 kb). **(g)** Retention of plasmids containing random genomic inserts, the EBV tethering sequence alone, or the entire EBV origin (containing both tethering and replicative sequences) compared to pUC19 in GM12878 cells (three biological replicates). Fold changes were computed using plasmid levels at day 14 post-transfection, normalizing to levels at day 2 to adjust for differential transfection efficiency across conditions. *P*-values computed by one-sided t-test.

**Extended Data Figure 3.**
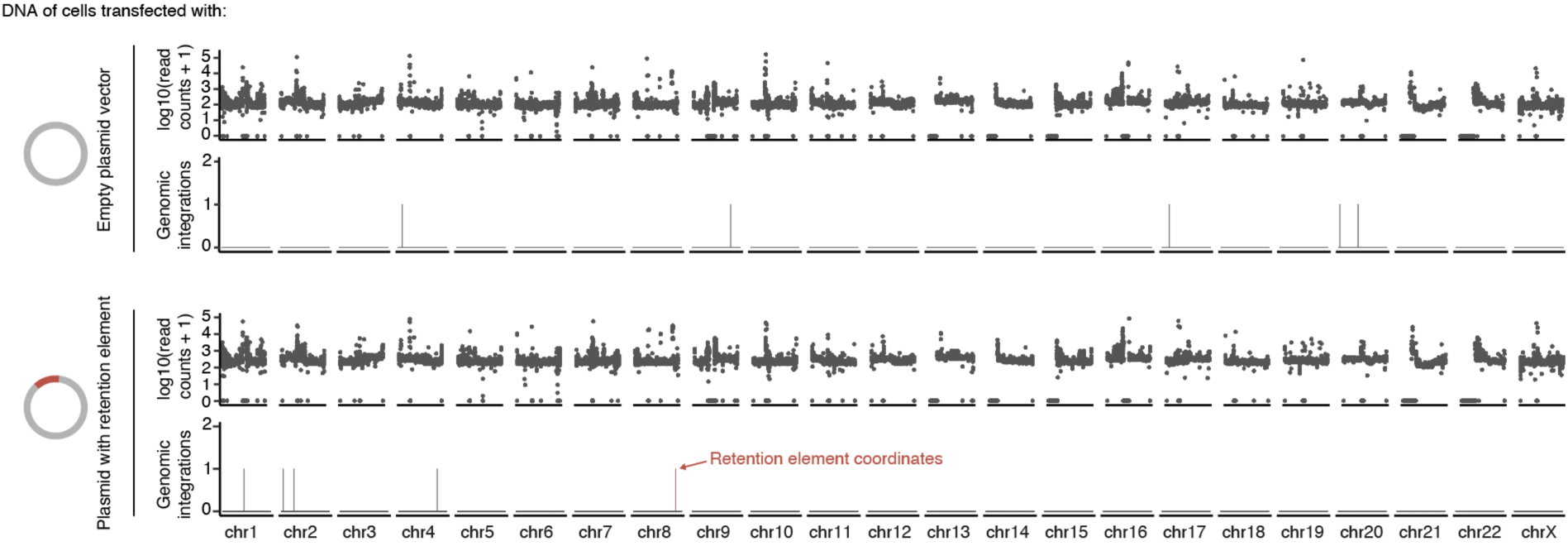
Chromosomal integration events of transfected plasmids containing a retention element are stochastic and occur at near-background levels. Genome-wide read coverage (non-overlapping 50 kb bins) and detection of chromosomal integration events (events per bin) of transfected plasmids in single-molecule long-read nanopore sequencing from cells transfected with either an empty plasmid vector (pUC19; top) or plasmid containing a retention element (pUC19_RE-C; bottom).

**Extended Data Figure 4.**
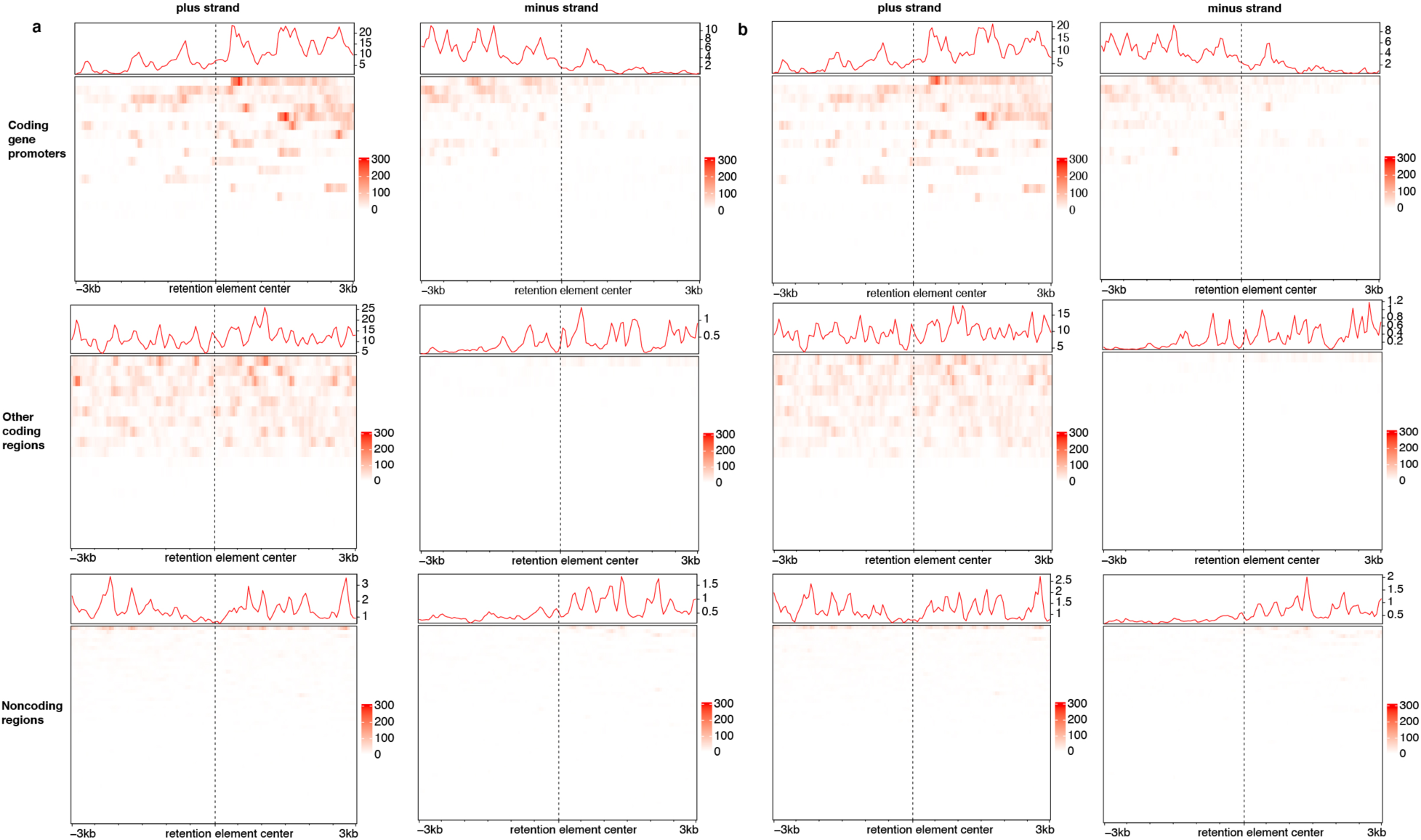
Many, but not all retention elements represent sites of active nascent transcription. **(a)** Histograms and heatmaps of COLO320DM GRO-seq signal from biological replicate 1, computed over 50 bp bins within 3 kb of the midpoints of retention elements located within the genomic coordinates of the COLO320DM ecDNA. Retention elements were divided into 3 categories based on overlap with genomic annotations: those that overlap with coding gene promoters, other portions of coding genes, or noncoding regions. X-axis directionality is consistent for both strands. **(b)** Heatmap of COLO320DM GRO-seq signal from biological replicate 2 within 3 kb of the midpoints of retention elements located within the genomic coordinates of the COLO320DM ecDNA.

**Extended Data Figure 5.**
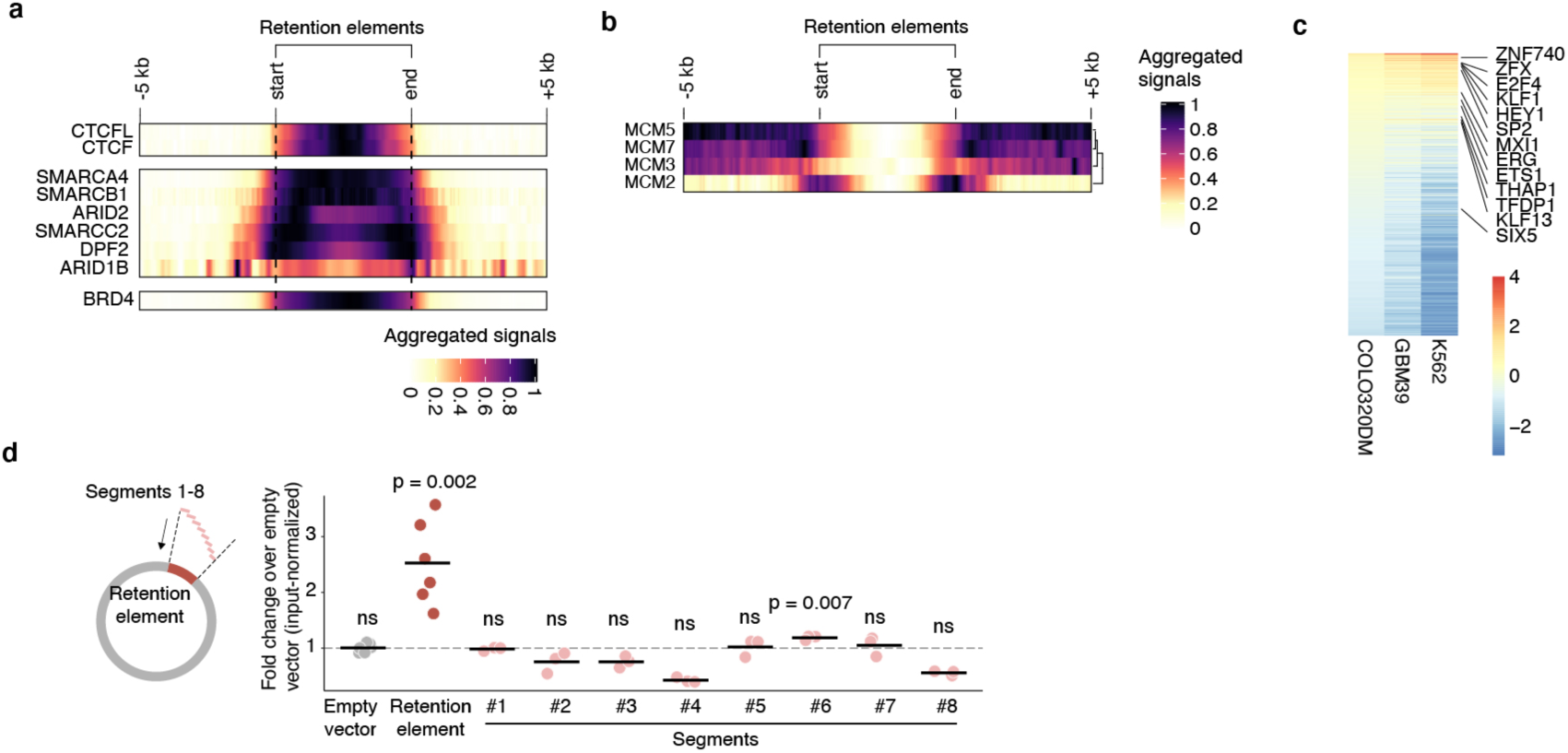
Additional sequence features of retention elements. **(a)** ENCODE ChIP-seq signals of the indicated proteins in K562 cells surrounding retention elements identified in the same cell line. **(b)** ENCODE ChIP-seq signals of components of the replication licensing complex in K562 cells surrounding retention elements identified in the same cell line. **(c)** Motif enrichment (log2 fold change) of transcription factor motifs in retention element intervals identified in COLO320DM, GBM39, and K562 cells relative to random genomic intervals. **(d)** Episomal retention of plasmids containing 8 overlapping 500-bp tiles of a retention element (RE-C) in COLO320DM cells measured by quantitative PCR (six biological replicates for empty vector and retention element conditions, three for others). *P*-values computed by one-sided t-test.

**Extended Data Figure 6.**
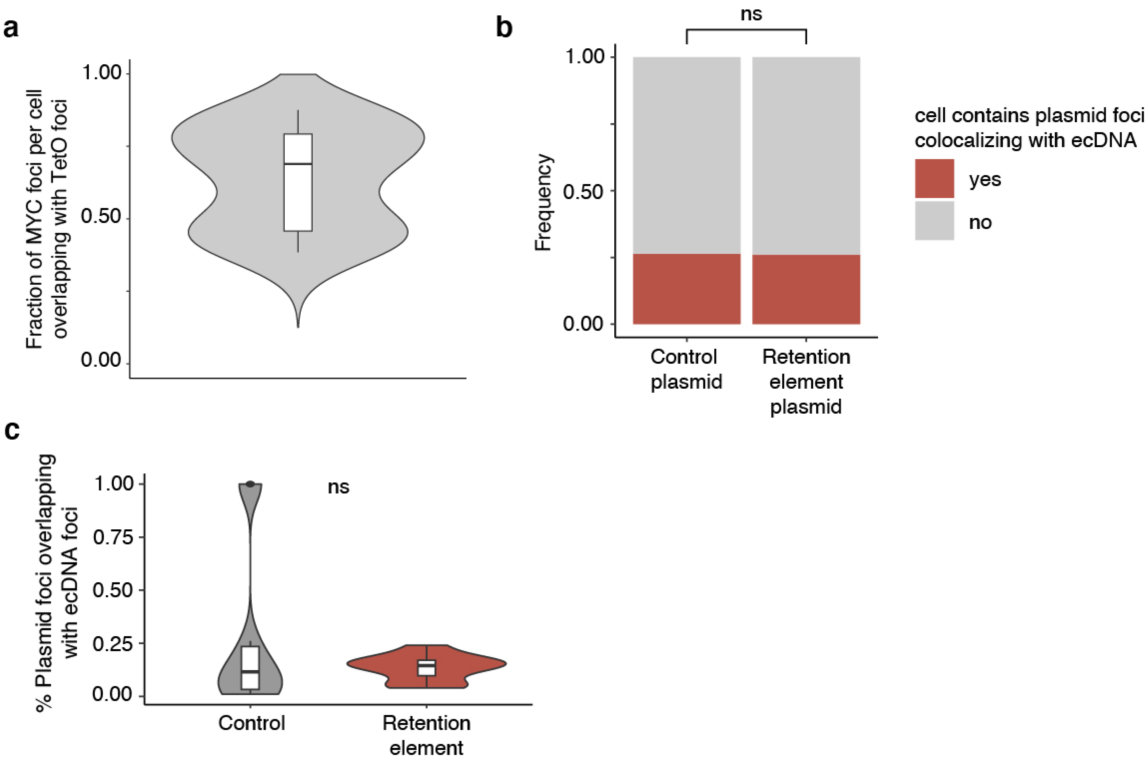
Summary of COLO320DM live cell imaging line. **(a)** Fraction of *MYC* ecDNA foci with overlapping TetO foci for each metaphase cell, indicating the percentage of labeled ecDNAs per cell (*n* = 20 cells). Box plot parameters as in Fig. 2. **(b)** Frequency of cells containing plasmid foci (either control or retention element plasmids) that colocalize with TetO-labeled ecDNA foci. *n* = 38 (control) and *n* = 46 (retention element) cells. *P*-value determined by one-sided hypergeometric test. **(c)** Percentages of plasmid foci area (either control or retention element plasmids) that colocalize with TetO-labeled ecDNA foci. *n* = 10 (control) and *n* = 12 (retention element) cells; only the subset of cells with plasmid foci that at least partially overlap with ecDNA foci are plotted here. Box plot parameters as in Fig. 2. *P-*value computed using a two-sample Wilcoxon test.

**Extended Data Figure 7.**
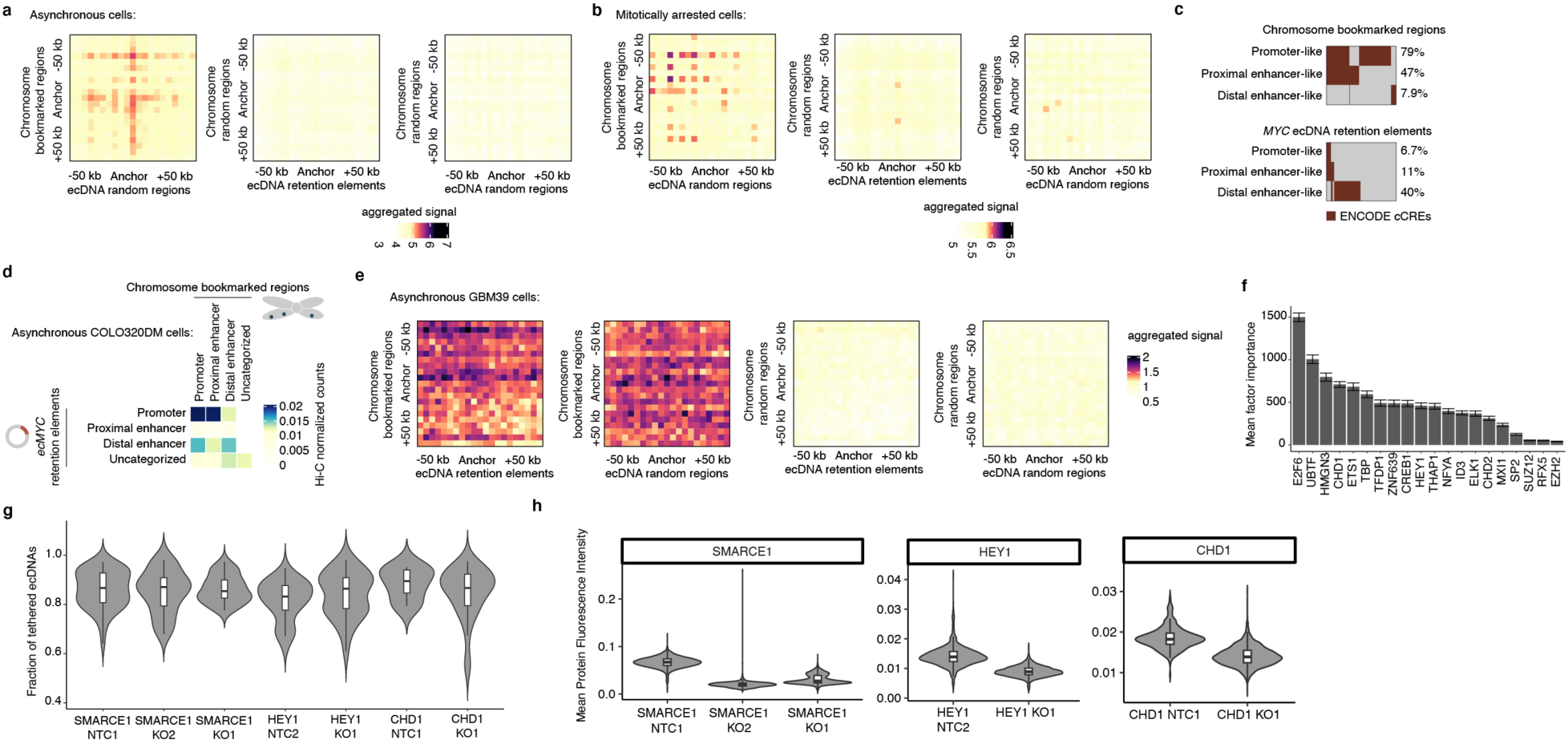
Chromatin interactions and functional annotations of chromosome bookmarked regions and *ecMYC* retention elements. (a-b) Aggregated peak analysis (APA) of Hi-C data of asynchronous (a) and mitotically arrested (b) COLO320DM cells. Heatmaps are summed percentile matrices of pairwise interactions between previously reported chromosome bookmarked regions (Methods) and a combined set of retention elements identified on the *MYC* ecDNA with 5-kb resolution, in which the chromosome bookmarked regions and/or the *ecMYC* retention elements are randomized. **(c)** Chromosome bookmarked regions or *ecMYC* retention elements with the indicated ENCODE cCRE annotations. **(d)** Hi-C heatmap of pairwise interactions between the *MYC* ecDNA retention elements and chromosome bookmarked regions with the indicated ENCODE cCRE annotations in asynchronous cells. Hi-C counts are normalized to number of interactions as well as bin sizes. **(e)** APA of Hi-C data of asynchronous GBM39 cells. **(f)** Importance scores (error bars show s.e.m.) indicating the relative contribution of each bookmarking factor to the cumulative distribution of retention elements. Scores represent the mean incremental number of retention elements containing binding sites for each factor over 1000 randomized cumulative distributions of the 20 bookmarking factors shown. Bookmarking factors are displayed in order of ChIP-seq peak enrichment within retention elements relative to random genomic intervals. **(g)** Fraction of tethered ecDNAs following CRISPR/Cas9 knockouts of selected bookmarking factors in mitotic COLO320DM cells. Box plot parameters as in Fig. 2. *n* = 55 (SMARCE1 NTC1), *n* = 42 (SMARCE1 KO1), *n* = 39 (SMARCE1 KO2), *n* = 34 (HEY1 NTC2), *n* = 33 (HEY1 KO1), *n* = 8 (CHD1 NTC1), *n* = 36 (CHD1 KO1) cells. **(h)** Mean immunofluorescence intensity of selected bookmarking factors in cells receiving targeting guide RNAs or non-targeting control (NTC) guides. *n* = 1874 (SMARCE1 NTC1), *n* = 2217 (SMARCE1 KO1), *n* = 1371 (SMARCE1 KO2), *n* = 1459 (HEY1 NTC2), *n* = 1976 (HEY1 KO1), *n* = 316 (CHD1 NTC1), *n* = 2730 (CHD1 KO1) cells. Box plot parameters as in Fig. 2.

**Extended Data Figure 8.**
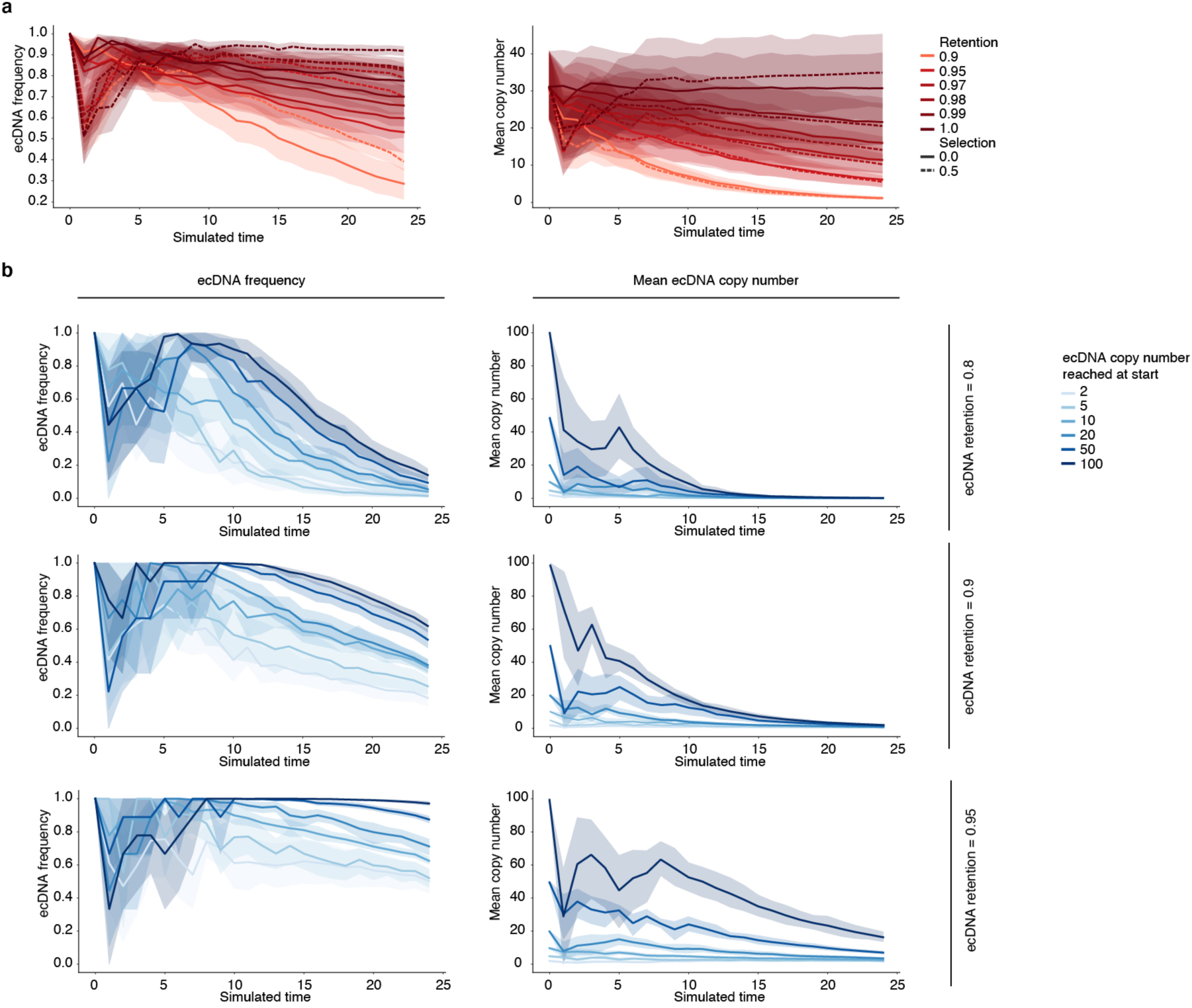
Evolutionary modeling of ecDNA retention and selection in growing cancer cell populations. **(a)** Time-resolved simulated trajectories of ecDNA frequency and mean copy number (95% confidence intervals shaded) across 25 simulated time units with various selection and retention values. **(b)** Time-resolved simulated trajectories of ecDNA frequency and mean copy number (95% confidence intervals shaded) across 25 simulated time units stratified by the number of initial ecDNA copies present in the parental cell. Trajectories are reported for various levels of retention. Selection is fixed at 0.5.

**Extended Data Figure 9.**
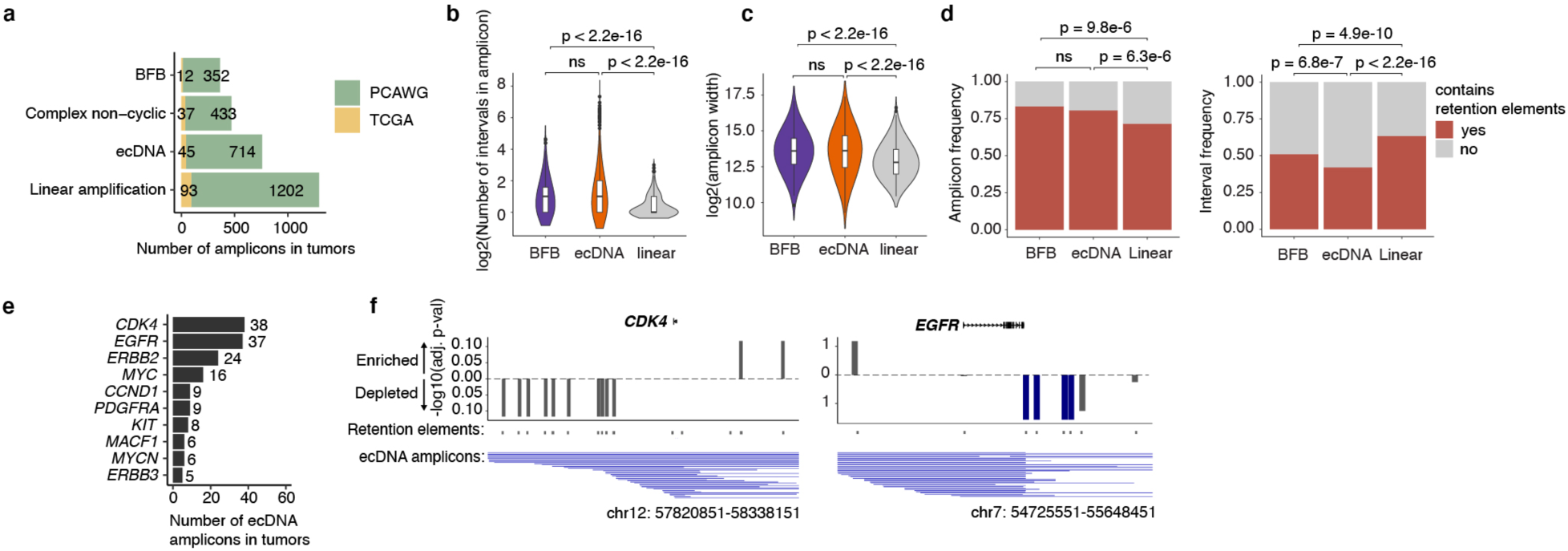
Summary statistics of DNA amplifications identified in WGS data of patient tumor samples. **(a)** Patient samples analyzed and classification of amplicons identified. **(b)** Number of genomic intervals implicated in each amplicon (i.e., degree of genomic rearrangement within an amplicon) across amplicon classes. *n* = 364 (BFB), *n* = 759 (ecDNA), and *n* = 1295 (linear) amplicons. Box plot parameters as in Fig. 2. *P*-values computed using two-sample Wilcoxon tests. **(c)** Amplicon widths (in bp) across amplicon classes. *n* = 364 (BFB), *n* = 759 (ecDNA), and *n* = 1295 (linear) amplicons. Box plot parameters as in Fig. 2. *P*-values computed using two-sample Wilcoxon tests. **(d)** Frequency of amplicons (left) or amplicon intervals (segments; right) containing at least one retention element across classes. *P-*values determined by one-sided hypergeometric test. **(e)** Top 10 oncogenes most frequently amplified as ecDNAs in analyzed patient samples. **(f)** Frequency of co-amplification of *CDK4* (left) or *EGFR* (right) with neighboring retention elements (within 250 kb of gene midpoint) in observed ecDNA amplicons (below each plot) reconstructed from patient samples relative to corresponding oncogene-containing random genomic intervals drawn from an equivalent size distribution.

**Extended Data Figure 10.**
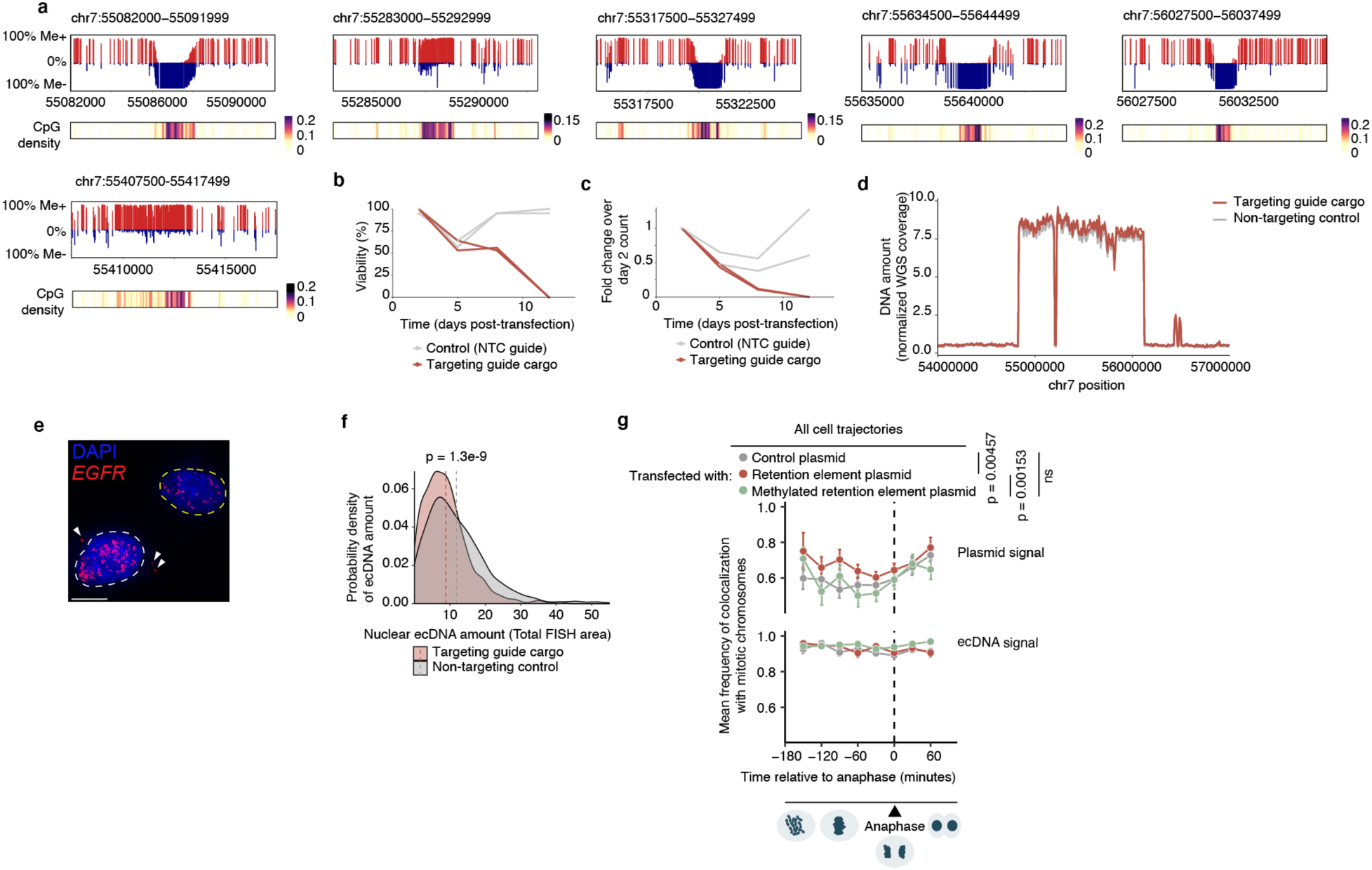
Hypomethylated CpG state is essential to retention element function. **(a)** 5mC methylation status of individual CpG sites and their density within and surrounding retention elements on the *EGFR* ecDNA in GBM39 cells as measured in single-molecule long-read nanopore sequencing. **(b)** Viability of cells expressing CRISPRoff and a targeting guide cargo or non-targeting control over time. Cells were sorted at day 2 post-transfection and tracked until day 12, when no live targeted cells remained. Each line represents an independent biological replicate. **(c)** Counts of cells expressing CRISPRoff and a targeting guide cargo or non-targeting control guide RNA over time. Cells were sorted at day 2 post-transfection and tracked until day 12, when no live targeted cells remained. Each line represents an independent biological replicate. **(d)** Abundance of ecDNA following CpG methylation of retention elements by CRISPRoff at 5 days post-transfection compared to cells expressing a non-targeting control guide RNA in WGS coverage. (**e**) Representative image showing ecDNA foci lost from the nucleus in an interphase GBM39 cell 5 days after transfection with CRISPRoff and a guide cargo targeting retention elements (*n* = 50 image positions). Scale bar, 10 µm. **(f)** Abundance of nuclear ecDNA measured by nuclear *EGFR* DNA FISH signal at 5 days after transfection of CRISPRoff and guide cargo targeting retention elements compared to cells expressing a non-targeting control guide RNA*. P*-value computed using two-sided two-sample Kolmogorov-Smirnov test. **(g)** Mean cell trajectories of methylated retention element plasmid (*n* = 51 cells) or *ecMYC* DNA signal colocalization with chromosomes throughout mitosis. Mean cell trajectories include all time points with more than 3 cells. Measurements for the control and unmethylated retention element plasmid conditions are reproduced from Figure 3d. Error bars show s.e.m. *P*-values determined by two-sided paired t-test of the means.

## METHODS

### Cell culture

The GBM39 neurosphere cell line was derived from a patient with glioblastoma undergoing surgery at Mayo Clinic, Rochester, Minnesota as described previously^61^. The COLO320DM and K562 cell lines were purchased from ATCC, and the GM12878 cell line was purchased from the Coriell Institute for Medical Research. The colorectal cancer cell line COLO320DM and the immortalized chronic myelogenous leukemia cell line K562 were cultured in Roswell Park Memorial Institute (RPMI) 1640 with GlutaMAX (Thermo Fisher Scientific, 61870127) supplemented with 10% FBS (Thermo Fisher Scientific, A3840002) and 1% Penicillin-Streptomycin (Thermo Fisher Scientific, 15140163). GBM39 cells were maintained in DMEM/F12 (Thermo Fisher Scientific, 11320082), B-27 supplement (Thermo Fisher Scientific, 17504044), 1% Penicillin-Streptomycin, human epidermal growth factor (EGF, 20ng/ml; Peprotech, AF-100-15), human fibroblast growth factor (FGF, 20ng/ml; Peprotech, AF-100-18B), and heparin (5ug/ml; Sigma-Aldrich, H3149). The lymphoblastoid cell line GM12878 was grown in RPMI 1640 with GlutaMAX supplemented with 15% FBS and 1% Penicillin-Streptomycin. The COLO320DM live cell imaging line was cultured in DMEM (Corning, 10-013-CV) supplemented with 10% FBS and 1% Penicillin-Streptomycin-Glutamine (Thermo Fisher Scientific, 10378016). GBM39 neurospheres were previously authenticated by the Mischel lab using metaphase DNA FISH^12^; other cell lines obtained from ATCC and Coriell were not authenticated. All cell lines tested negative for mycoplasma contamination.

### Analysis of ecDNA hitchhiking in IF-DNA-FISH of anaphase cells

Analysis of ecDNA hitchhiking in IF-DNA-FISH of anaphase cells was performed on raw images used in a previous publication^5^. Mitotic cells were identified using Aurora kinase B, which identifies daughter cell pairs undergoing mitosis, as previously described^5,6^. Colocalization analysis for ecDNAs with mitotic chromosomes in GBM39 cells (*EGFR* ecDNA), PC3 cells (*MYC* ecDNA), SNU16 cells (*FGFR2* and *MYC* ecDNAs) and COLO320DM cells (*MYC* ecDNA) described in **Figure 1** was performed using Fiji (v.2.1.0/1.53c)^62^. Images were split into the FISH color + DAPI channels, and signal threshold set manually to remove background fluorescence. DAPI was used to mark mitotic chromosomes; FISH signals overlapping with mitotic chromosomes were segmented using watershed segmentation. Colocalization was quantified using the ImageJ-Colocalization Threshold program and individual and colocalized FISH signals in dividing daughter cells were counted using particle analysis.

### Retain-seq

We cloned random genomic sequences into the pUC19 plasmid backbone for the Retain-seq experiments. pUC19 is a simple, small (∼2.7 kb) vector that lacks a mammalian origin of replication and contains few sequences that could be immunogenic or have mammalian promoter or enhancer activity; thus, we believe pUC19 represents an inert and selectively neutral backbone. Therefore, changes in plasmid persistence can be more confidently ascribed to insert sequences as opposed to backbone components under selection. To generate a pool of random genomic sequences, we first fragmented the genomic DNA of GM12878 cells via transposition with Tn5 transposase produced as previously described^63^, in a 50-µl reaction with TD buffer^64^, 50 ng DNA and 1 µl transposase. The reaction was performed at 37°C for 5 minutes, and transposed DNA was purified using MinElute PCR Purification Kit (Qiagen, 28006). GM12878 human B lymphoblastoid cells were selected as the genome of origin due to their relatively low copy-number variability and the presence of an EBV genome as a positive control; the majority of inserts ranged from 600-1300bp. The resulting mixture of genomic DNA fragments was then amplified using 500 nM forward (p5_pUC19_SmaI_20bp) and reverse (p7_pUC19_SmaI_20bp) primers using NEBNext High-Fidelity 2× PCR Master Mix (NEB, M0541L) followed by gel purification of DNA fragments between 400 bp and 1.5 kb. To insert the mixture of genomic DNA fragments into a plasmid, the pUC19 vector (Invitrogen) was linearized with SmaI, purified using NucleoSpin® Gel and PCR Clean-up (Macherey-Nagel, 740609.250) and the genomic fragments were inserted into the backbone using Gibson assembly (New England Biolabs, NEB). The DNA product was electroporated into Endura Competent Cells (Biosearch Technologies, 60242-2) using a MicroPulser Electroporator (Bio-Rad; default bacteria setting) following the manufacturer’s protocol, and the resulting mixed episome library was prepared using the HiSpeed Plasmid Maxi Kit (Qiagen, 12663). The analysis of representation of DNA sequences in this mixed episome library as well as retained episomes in transfected cells is described below.

COLO320DM and K562 cells were seeded into a 15cm dish per biological replicate at a density of 1 × 10^7^ cells in 25 ml of media; GBM39 cells were seeded into a T75 flask at a density of 5 × 10^6^ cells in 25 ml of media. Each cell line was incubated overnight. COLO320DM, GBM39, and K562 cells were transfected with 15 µg input mixed episome library using the Lipofectamine 3000 Transfection Reagent following the manufacturer’s directions. 1.5 × 10^7^ GM12878 cells were electroporated with 50 µg input mixed episome library using the Neon Transfection System (Thermo Fisher Scientific, MPK5000). Briefly, the cells were counted, centrifuged at 300g for 5 min, and washed twice with PBS before resuspension in Neon Resuspension Buffer to a density of 4.2 × 10^6^ in 70 µl of buffer; input mixed episome library was also diluted to a density of 14 µg in 70 µl with Neon Resuspension Buffer. 70 µl of cell suspension and 70 µl of library were mixed and electroporated according to the manufacturer’s instructions using a 100 µl Neon pipet tip under the following settings (1200 V, 20 ms, 3 pulses). 5 electroporation reactions were pooled per replicate of GM12878 Retain-seq screens.

Cells were incubated for 2 days prior to the first subculture to permit recovery from transfection, and then sub-cultured every 3-4 days afterward as dictated by each cell line’s doubling time. Once each cell line reached a count of 100-400 million cells per replicate, we harvested all but 10 million cells, which were maintained in culture and passaged in the same manner until all subsequent time points had been collected (for a maximum of 3 time points per cell line). Thus, COLO320DM cells were harvested at days 7, 14, and 21 following transfection with a total cell count of approximately 4 × 10^8^ cells at each time point, per replicate; GBM39 was harvested at days 10, 20, and 30 with total cell counts of approximately 1.5 × 10^8^ per replicate; K562 was harvested at days 6, 12, and 18 with cell counts of approximately 4.5 × 10^8^ per replicate; and GM12878 was harvested at day 12 with a cell count of approximately 2 × 10^8^.

The output plasmid library was extracted using the HiSpeed Plasmid Maxi Kit (Qiagen, 12663) and concentrated to a final volume of 50 µl by isopropanol precipitation. DNA was precipitated with a 1:10 volume of 3M sodium acetate and 2 volumes of isopropanol, chilled at 4°C for 10 min, and centrifuged at 15,000*g* for 15 min at 4°C. The pellet was washed with 500 µl ice cold 70% ethanol and dissolved in 50 µl Buffer EB (Qiagen, 19086).

To enrich for input mixed episome library inserts, a preliminary PCR amplification (PCR1) of 10 cycles using primers (at 500 nM) annealing to the pUC19 vector (forward: pUC19_SmaI_5prime_fwr; reverse: pUC19_SmaI_3prime_rev) were performed on the concentrated DNA using NEBNext High-Fidelity 2x PCR Master Mix (NEB, M0541L). Each PCR1 reaction used a maximum of 2 µg concentrated DNA as template, with reactions assembled successively until all concentrated DNA was consumed; all reactions for a given sample were pooled following PCR1 and purified using the NucleoSpin Gel & PCR Clean-up kit (Macherey-Nagel, 740611), resulting in PCR product 1. Due to variability in insert size and the amount of retained plasmid DNA in the output library, artificial overrepresentation of fragments caused by PCR over-cycling represented a concern for subsequent sequencing. Thus, we used quantitative PCR to identify the cycle before saturation and halted amplification at this point. For quantitative PCR, 50 ng of DNA from PCR product 1, NEBNext High-Fidelity 2x PCR Master Mix, 500 nM forward and reverse primers (forward: p5_adapter_only; reverse: p7_adapter_only), and 1 µl of 25x SYBR Green I (diluted from 10,000x stock; Thermo Fisher Scientific, S7563) were used in a 50 µl reaction. SYBR Green signal of amplification products was measured in technical triplicates per reaction using Lightcycler 480 (Roche) and plotted against cycle number to identify the PCR cycle before saturation. According to the cycle numbers identified by this quantitative PCR step, we then performed PCR2 by amplifying PCR product 1 (50 ng DNA) using the same primers as in the quantitative PCR: 5, 10, and 12 PCR cycles for days 7, 14, and 21 of the COLO320DM experiment; 5, 11, and 18 PCR cycles for days 10, 20, and 30 of the GBM39 experiment; 5, 11, and 17 PCR cycles for days 6, 12, and 18 of the K562 experiment; and 10 PCR cycles for day 12 of the GM12878 experiment. We also collected a day 17 time point from the GM12878 experiment (amplified using 16 PCR cycles) that was specifically used to study retention of the EBV FR element, as this time point was assumed to be more comparable to the second time point in other cell lines. Next, output DNA from this step (PCR product 2) was purified using the MinElute PCR Purification Kit (Qiagen, 28006) and then transposed with Tn5 transposase produced as previously described^63^ in a 50 µl reaction with TD buffer^64^, 50 ng DNA (PCR product 2), and 1 µl transposase. The reaction was performed at 50°C for 5 min, and transposed DNA was purified using the MinElute PCR Purification Kit (Qiagen, 28006). The above PCR steps and transposition were also carried out on the input mixed episome library originally used for cell transfection but with 25 ng of input mixed episome library for PCR1. According to the cycle numbers identified by this quantitative PCR step, we then amplified PCR product 1 (1 ng DNA) over 9 PCR cycles (PCR2). Finally, the prior PCR steps and transposition were also performed on a dilution series of 10 ng, 1 ng, 0.1 ng, and 0.01 ng of input mixed episome library as PCR1 template DNA in order to standardize analysis of screen output across varying DNA amounts.

Sequencing libraries were generated by 5 rounds of PCR amplification on the transposed PCR product 2 using NEBNext High-Fidelity 2× PCR Master Mix (NEB, M0541L) with primers bearing i5 and i7 indices, purified using the SPRIselect reagent kit (Beckman Coulter, B23317) with left-sided size selection (1.2x), and quantified using Agilent Bioanalyzer 2100. Libraries were diluted to 4 nM and sequenced on the Illumina NovaSeq 6000 platform.

Primer sequences are listed in **Supplementary Table 2**.

### Retain-seq analysis

Sequenced episome library reads were trimmed of adapter content with Trimmomatic^65^ (version 0.39), aligned to the hg19 genome using BWA MEM^66^ (0.7.17-r1188), and PCR duplicates removed using Picard’s MarkDuplicates (version 2.25.3). Read counts were then obtained for 1-kilobase windows across the reference hg19 genome using bedtools (v.2.30.0). Windows with fewer than 10 reads within 1 kb in the input episome library were filtered out.

Next, read counts were normalized to total reads and scaled to counts per million (CPMs). We filtered out blacklist regions of the genome^67^ and windows with extreme outlying read counts in the input episome library (more than three standard deviations above the mean read count). To determine how genome coverage is affected by input DNA amount, we measured read counts of 1-kb genomic bins from sequencing of serial dilutions of the input episome library. Based on this serial dilution experiment which showed consistent representation of DNA sequences down to 0.1 ng of input DNA, at which the genome representation was nearly identical to 1 ng and 10 ng of input DNA in the top 50% of genomic bins (**Extended Data Figure 1b**; 0.01 ng showed substantial library dropout and signs of skewing), we focused our subsequent analysis of Retain-seq on time points at which at least 50% of genomic bins are represented (i.e. above 10 reads within a 1-kb window. GBM39 at day 30 showed low genome representation and was excluded from subsequent analysis. K562 at day 18 showed a large drop in genome representation and was excluded from subsequent analysis; **Extended Data Figure 2a**).

We then calculated the log2 fold change of each genomic window in each sample over the input episome library by dividing the respective CPMs followed by log-transformation. Regions of the background genome with copy-number amplification in the cells retaining the episome library can elevate the background sequencing reads aligning to those regions. To remove such background genomic noise, we calculated the median log2 fold change values of the neighboring windows +/- 5 kb from each 1-kb window and normalized the log2 fold change of each 1-kb window to its corresponding neighbor average. Thus, any enriched episome sequence was required to have increased signal both compared to the input level as well as its neighboring sequences in its position in the reference human genome. Z scores were calculated using the formula z = (x-m)/S.D., where x is the log2 fold change of each 1-kb window, m is the mean log2 fold change of the sample, and S.D. is the standard deviation of the log2 fold change of the sample. Z scores were used to compute upper-tail P values using the normal distribution function, which were adjusted with p.adjust in R (v.3.6.1) using the Benjamini-Hochberg Procedure to produce false discovery rate (FDR) values. To identify episomes enriched in various cell lines, we identified 1-kb windows with FDR < 0.1 in two biological replicates at any of the time points for sample collection.

### Plasmid cloning

Retention element individual validations: pUC19 (empty vector) was digested with SmaI; each retention element sequence (RE-A: chr7:55321959-55323480; RE-B: chr7:55432848-55434854; RE-C: chr8:127725819-127727938; RE-D: chr7:56032209-56033389; RE-E: chr7:55086476-55088263; RE-F: chr7:55639062-55640378) was PCR amplified via a 2-step nested PCR from genomic DNA derived from the GM12878 cell line and inserted into the empty vector by Gibson assembly using the NEBuilder HiFi 2x DNA Assembly Master Mix (NEB, E2621L) in accordance with the manufacturer’s protocol. The resulting plasmids were named: pUC19_RE-A, pUC19_RE-B, pUC19_RE-C, pUC19_RE-D, pUC19_RE-E, and pUC19_RE-F.

To clone pUC19 plasmids containing the EBV tether (pUC19_FR) or the entire viral origin (tether and replicator; pUC19_oriP), the viral tether (FR element; EBV:7421-8042) and viral origin (oriP; EBV:7338-9312) sequences were PCR-amplified using the pHCAG-L2EOP plasmid (Addgene, 51783)^68^ as a template and inserted into SmaI-digested pUC19 by Gibson assembly.

To clone pUC19 plasmids with 2 or 3 copies of a retention element (RE-C: chr8:127725819-127727938; pUC19_2RE and pUC19_3RE), we digested pUC19_RE-C with HindIII and inserted a second copy of the retention element (amplified by PCR primers pUC19_2RE forward and pUC19_2RE reverse) by Gibson assembly to generate pUC19_2RE. To generate pUC19_3RE (3 copies of the retention element), pUC19_2RE was digested with SacI and a third copy of the retention element (amplified by PCR primers pUC19_3RE forward and pUC19_3RE reverse) was inserted by Gibson assembly.

To clone the pUC19 plasmid containing the CMV promoter (pUC19_CMV), the CMV promoter was PCR-amplified (primers pUC19_CMV forward and pUC19_CMV reverse) using the pGL4.18 CMV-Luc plasmid (pGL4; Addgene, 100984)^69^ as a template and inserted into HindIII-digested pUC19 by Gibson assembly. To clone the pGL4 vector containing a retention element (RE-C: chr8:127725819-127727938; pGL4_RE-C), we digested pGL4 with MfeI and BamHI for the backbone and PCR-amplified the retention element sequence from GM12878 genomic DNA (primers pGL4_RE1 forward and pGL4_RE1 reverse). The PCR product was gel purified, digested with BsaI and BamHI, and ligated to the vector backbone using the DNA Ligation Kit Version 2.1 (Takara Bio, 6022) following the manufacturer’s protocol.

For cloning individual overlapping tiles of a retention element (RE-C: chr8:127725819-127727938; tiles were 500 bp each, with the first 250 bp overlapping with the previous tile and the latter 250 bp with the following tile), each tile was amplified by PCR using pUC19_RE-C as a template; pUC19 was digested with SmaI and each tile sequence was inserted by Gibson assembly.

The plasmids for live cell imaging were designed based on a previously published pGL4 vector for a dual luciferase assay^23^, which contains a retention element (chr8:128,804,981-128,806,980, hg19) overlapping with the *PVT1* promoter termed RE-G. To insert LacO repeats for imaging, we first inserted multiple enzyme sites (GTCGACTGTGCTCGAGAACACGGATCCTATGCTCGTACG) by Gibson assembly following digestion with BamHI. Next, the vector was digested with SalI and Bsiwi, and ligated with an array of 256 LacO copies that was obtained by digestion of a pLacO-ISce1 plasmid (Addgene, 58505)^70^ with SalI and Acc65I. To create a control plasmid that does not contain the retention element, the vector was digested with KpnI and BglII. The plasmid sequences are verified by Sanger sequencing. The LacO repeats in the plasmids were further verified with agarose gel due to its large size. All enzymes and Gibson assembly mix are purchased from NEB. All primer sequences are listed in **Supplementary Table 2**.

### Quantitative PCR analysis of plasmid retention

To assess the retention of individual plasmids transfected into cells, we seeded K562 or COLO320DM cells into 6-well plates at a density of 3 × 10^5^ cells in 3 ml of media per well and incubated overnight. The next morning, the cells were transfected with 0.5 µg plasmid per well using the Lipofectamine 3000 Transfection Reagent (Thermo Fisher Scientific) following the manufacturer’s protocol. 6 × 10^5^ GM12878 cells were electroporated with 2 µg plasmid per well using the Neon Transfection System. Cells were counted, centrifuged at 300g for 5 min, and washed twice with PBS before resuspension in Neon Resuspension Buffer to a density of 4.2 × 10^5^ in 7 µl of buffer; plasmid was also diluted to a density of 1.4 µg in 7 µl with Neon Resuspension Buffer. 7 µl of cell suspension and 7 µl of plasmid were mixed and electroporated according to the manufacturer’s instructions using a 10 µl Neon pipet tip under the following settings (1200 V, 20 ms, 3 pulses). 2 electroporation reactions were pooled per replicate and plated into a 12-well plate in 1.5 ml of media per well. Cell cultures were split every 2-4 days and fresh media was added. To quantify plasmid DNA in cells at various time points, genomic DNA was extracted from cells using the DNeasy Blood & Tissue Kit (Qiagen, 69504). Quantitative PCR was performed in technical duplicates using 50-100 ng of genomic DNA, 2x LightCycler 480 SYBR Green I Master mix (Roche, 04887352001), and 125 nM forward and reverse primers (primers pUC19_F and pUC19_R, annealing to the pUC19 vector backbone; for plasmids with the pGL4 vector backbone, primers pGL4_F and pGL4_R were used). Relative plasmid DNA levels were calculated by normalizing to GAPDH controls (primers GAPDH_F and GAPDH_R). DNA levels were further normalized to the day 2 levels to account for variability in transfection efficiencies and to cells transfected with an empty plasmid vector control. *P*-values were calculated in R using a Student’s *t*-test by comparing the relative fold change of biological replicates at various time points with respect to the input levels at day 2. Primer sequences are listed in **Supplementary Table 2**.

### Analysis of potential genomic integration of plasmids

COLO320DM cells were seeded into two wells of a 6-well plate, transfected with 0.5 µg of pUC19 or pUC19_RE-C per well, and passaged as described previously in the section “Quantitative PCR analysis of plasmid retention.” At day 8, high-molecular-weight genomic DNA was extracted from cells with the Puregene Cell Core Kit (Qiagen, 158046) and long-read sequencing libraries were prepared using the Ligation Sequencing Kit V14 (Oxford Nanopore Technologies, SQK-LSK114) in accordance with the manufacturer’s protocol. Libraries were loaded onto R10.4.1 flow cells (Oxford Nanopore Technologies, FLO-PRO114M) and sequenced on the PromethION platform (Oxford Nanopore Technologies). Basecalling from raw POD5 data was performed using the High accuracy (HAC) DNA model in Dorado (Oxford Nanopore Technologies, version 0.5.2). Fastq files were generated using samtools bam2fq (version 1.6)^71^, aligned to a custom reference (hg19_pUC19) comprising the pUC19 sequence appended to the hg19 genome using minimap2 (version 2.17)^72^, and sorted and indexed using samtools; alignments shorter than 1 kb and with mapping quality below 60 were discarded. Structural variants were then called using Sniffles (version 2.2)^73^ using the hg19_pUC19 reference and the following parameters: “--allow-overwrite --output-rnames --non-germline --long-ins-length 3000”. Integration events were identified from Sniffles output (.vcf) as Breakends (Translocations) between the pUC19 sequence and chromosomes.

### ENCODE data integration

To perform meta-analysis of protein binding sites within retention elements, ENCODE data were downloaded in “bigWig” format using the files.txt file returned from the ENCODE portal (https://www.encodeproject.org) and the following command: “xargs -n 1 curl -O -L < files.txt”. K562 retention element coordinates were converted from the h19 to hg38 build using the UCSC LiftOver tool (R package liftOver, version 1.18.0). To plot heatmaps of protein binding within retention elements, we used the “computeMatrix” function in deepTools (version 3.5.1) using the “scale-regions” mode, specified each “bigWig” file using “--scoreFileName”, and a .bed file containing hg38 retention element coordinates using “--regionsFileName”, along with the following parameters: “-- regionBodyLength 5000 --beforeRegionStartLength 5000 --afterRegionStartLength 5000 --binSize 20 –skipZeros”. Each resulting matrix was aggregated by computing column means using the colMeans function in R and rescaled to 0-1 using the “rescale” function in the scales (version 1.3.0) package in R.

To analyze overlap of various genomic annotation classes within retention elements, coordinates of each genomic annotation type were first obtained using the R packages TxDb.Hsapiens.UCSC.hg19.knownGene (genes; version 3.2.2) and TxDb.Hsapiens.UCSC.hg19.lincRNAsTranscripts (lncRNAs; version 3.22). “All promoters” comprised sequence 1500 bp upstream to 200 bp downstream from the transcription start site for all transcripts in the TxDb objects, extracted using the “promoters” function. 5’ UTR, 3’ UTR, intron, and exon sequences were extracted using the “fiveUTRsByTranscript”, “threeUTRsByTranscript”, “intronicParts”, and “exonicParts” functions respectively while coding and lncRNA promoters were each subsets of the total promoters list. Downstream intergenic regions represent non-genic sequences within 1500 bp of each transcription termination site while distal intergenic regions were classified as non-genic sequences beyond 1500 bp of the TSS and 1500 bp of the TTS; coordinates were computed using the “flank” and “setdiff” functions in the R package GenomicRanges (version 1.46.1).

To analyze enrichment of transcription factor binding sites within retention elements, uniformly processed transcription factor ChIP-seq data (aligned to the hg38 genome) from the K562 cell line were downloaded as a batch from the Cistrome Data Browser (Cistrome DB)^74^. Datasets that failed to meet more than one of the following quality thresholds were excluded: raw sequence median quality score (FastQC score) ≥ 25; ratio of uniquely mapped reads ≥ 0.6; PBC score ≥ 80%; union DNase I hypersensitive site overlap of the 5,000 most significant peaks ≥ 70%; number of peaks with fold change above 10 ≥ 500; and fraction of reads in peaks ≥ 1%. Individual ChIP-seq datasets were imported as GenomicRanges (version 1.46.1) objects from narrowPeak or broadPeak files. For transcription factors with multiple ChIP-seq datasets, datasets were aggregated into a union peak set for subsequent analyses. To identify transcription factors that are enriched for binding within retention elements relative to random genomic intervals, a fold change was computed for each transcription factor comparing the percentage of retention element intervals overlapping with at least 1 transcription factor ChIP-seq peak (> 50% peak coverage) against the percentage of overlapping 1 kb genomic bins; *p*-values were computed in R (function “phyper”) using a hypergeometric test for over-representation and adjusted for multiple comparisons by the Bonferroni correction.

### Origins of replication overlap

Coordinates (in the hg19 reference) of origins of replication identified in the K562 cell line across 5 replicates of SNS-seq were published with Picard et al. and deposited in NCBI Gene Expression Omnibus (GEO) under accession GSE46189^75^. Retention elements or 1 kb genomic bins were considered overlapping if an origin of replication covered at least 25% of the queried interval (calculated in R using the package GenomicRanges, version 1.46.1). The enrichment *p-*value was computed in R using a hypergeometric test for over-representation.

### GRO-seq analysis

GRO-seq data of COLO320DM were published with Tang et al. and deposited in NCBI GEO under accessions GSM7956899 (replicate 1) and GSM7956900 (replicate 2)^76^. The subset of retention element coordinates from the COLO320DM, GBM39, or K562 cell lines located within the amplified intervals of the COLO320DM ecDNA was divided into three categories based on overlap with genomic annotations: 1) retention elements located entirely within coding gene promoters (within 2 kb of a coding gene TSS); 2) retention elements located elsewhere within the limits of coding genes; and 3) retention elements located within noncoding regions. Coordinates of these retention elements were then converted from the hg19 to hg38 build using the UCSC liftOver package (version 1.18.0) in R. GRO-seq signal within 3 kb of the midpoint of each retention element was presented in separate heatmaps using the EnrichedHeatmap package (version 1.24.0) for each strand and for each retention element category.

### Motif enrichment

A curated collection of human motifs from the CIS-BP database^77^ (“human_pwms_v2” in the R package chromVARmotifs, version 0.2.0)^78^ was first matched to the set of 1 kb bins spanning the hg19 reference to identify all such intervals of the human genome containing instances of each motif. Enrichment of each motif within retention elements was then calculated as a log2(fold change) of the fraction of retention element intervals (identified by Retain-seq in each cell type) containing motif instances compared to all genomic intervals.

### Live-cell imaging

The live cell imaging cell line was engineered from COLO320DM cells obtained from ATCC, as described in a previous publication^6^. TetO ecDNAs are labeled with TetR-mNeonGreen. Based on overlap between MYC and TetO FISH foci in metaphase spreads, 50-80% of ecDNA molecules in a given cell were typically labeled (**Extended Data Figure 6a**). The cells were further infected with the LacR-mScarlet-NLS construct and sorted for mScarlet-positive cells to enable stable expression of LacR-mScarlet protein. These cells were then subjected to nucleofection of either the control plasmid with LacO repeats, the plasmid containing a retention element (RE-G) with LacO repeats, or the *in vitro* CpG methylated retention element (RE-G) plasmid with LacO repeats. Specifically, 1 μg of plasmid were nucleofected into 400,000 cells following the standard nucleofection protocol from Lonza (Nucleofection code: CM-138) to visualize plasmid signal. Cells were seeded onto poly-D-lysine (10 μg/mL; Sigma-Adrich #A-003-E) coated 96-well glass-bottom plates (Azenta Life Sciences MGB096-1-2-LG-L) immediately after nucleofection and were imaged two days later. FluoroBrite DMEM (Gibco, A1896701) supplemented with 10% FBS and 1X Glutamax, along with 1:200 Prolong live antifade reagent (Invitrogen, P36975), was replenished 30 minutes prior to time-lapse imaging. Cells were imaged on a top stage incubator (Okolab) fitted onto a Leica DMi8 widefield microscope with a 63x oil objective, with temperature (37°C), humidity and CO2 (5%) controlled throughout the imaging experiment. Z-stack images were acquired every 30 minutes for a total of 4 to 18 hours. The images were processed using Small Volume Computational Clearing before maximum intensity projections were made for all frames.

### Live-cell imaging analysis

Maximum intensity projections were exported as TIFF files from the .lif files using imageJ. To analyze colocalization of LacR-LacO-plasmid foci or TetR-TetO-*MYC* ecDNA foci with mitotic chromosomes during anaphase, images of cells entering anaphase and telophase were exported for mitotic cells that had showed at least five distinct plasmid foci at the beginning of mitosis. The exported images were split into the different color channels, and signal threshold set manually to remove background fluorescence using Fiji (version 2.1.0/1.53c)^62^. Fluorescence signals were segmented using watershed segmentation. H2B-emiRFP670 signal was used to mark the boundaries of mitotic chromosomes of dividing daughter cells. All color channels except H2B were stacked and ROIs were drawn manually to identify the two daughter cells, and a third ROI was drawn around the space occupied by the pair of dividing daughter cells. Next, the colour channels were split again and image pixel areas occupied by fluorescence signals were analyzed using particle analysis. Fractions of ecDNAs colocalizing with mitotic chromosomes were estimated by fractions of FISH pixels within the daughter cell chromosome ROIs.

To perform time-resolved DNA segregation analysis, TIFF files were analyzed on Aivia (v.12.0.0) by first segmenting the condensed chromatin (labelled by H2B-emiRFP670), TetR-TetO-*MYC* foci, and LacR-LacO-plasmid foci of the mitotic cell, using a trained pixel classifier recognizing each of the elements. Each segmented chromatin and focus of interest was then selected manually and output as an object. The relative distance of each focus to its corresponding segmented chromatin’s periphery was output using the Object Relation Tool, by setting the ‘TetR/PVT1’ object as primary set and its corresponding ‘Chromatin’ object as secondary set, under default settings. The resulting data were exported to R (v.3.6.1). TetR-TetO-*MYC* foci or LacR-LacO-plasmid foci with more than 75% overlapping area with the ‘Chromatin’ object were considered colocalized and their relative distances to their corresponding segmented chromatin were replaced with 0. For each dividing cell, the fractions of plasmid or ecDNA foci colocalizing with mitotic chromosomes were calculated.

### Hi-C

For mitotic Hi-C of COLO320DM cells, COLO320DM cells were seeded into a 6 cm dish at a density of 0.5 × 10^6^ cells in 8 ml of RPMI media (11875-119) containing 10% fetal bovine serum (Fisher Scientific, SH30396.03) and 1% penicillin-streptomycin (Gibco, 15140-122), the cells were incubated overnight. Nocodazole (M1404-10MG) was dissolved in DMSO and added directly to the cells in the media to reach a final concentration of 100 ng/μl (8 μl of 100 ng/ml nocodazole was added to 8 ml RPMI media). After 16 hours of nocodazole treatment, both suspension and adherent cells were harvested for Hi-C analysis and flow cytometry analysis for cell cycle staining using propidium iodide (Invitrogen, 00699050). Flow cytometry verified that the cell population consisted mainly of cells with 4n DNA content after mitotic arrest. For interphase Hi-C of GBM39 (GBM39ec) cells, GBM39 cells were cultured as described above (section “Cell culture”.

To perform each Hi-C experiment, ten million cells were fixed in 1% formaldehyde in aliquots of one million cells each for 10 minutes at room temperature and combined after fixation. We performed the Hi-C assay following a standard protocol to investigate chromatin interactions^79^. Hi-C libraries were sequenced on an Illumina HiSeq 4000 with paired-end 75 bp reads for mitotic Hi-C of COLO320DM and an Illumina NovaSeq 6000 with paired-end 150 bp reads for interphase Hi-C of GBM39^80^.

### Hi-C analysis

Paired-end Hi-C reads were aligned to hg19 genome with the Hi-C-Pro pipeline^81^. Pipeline was set to default and set to assign reads to DpnII restriction fragments and filter for valid pairs. The data was then binned to generate raw contact maps which then underwent ICE normalization to remove biases. Visualization was done using Juicebox (https://aidenlab.org/juicebox/). Hi-C data from asynchronous COLO320DM and GBM39 cells were generated and processed in the same way in parallel with the mitotically arrested cells; asynchronous COLO320DM cell data were separately published with Kraft et al. 2024 (*bioRxiv*) and deposited in NCBI Gene Expression Omnibus (GEO) under accessions GSM8523315 (replicate 1) and GSM8523316 (replicate 2)^82^.

To analyze chromatin interactions with retention elements on *ecMYC*, the combined set of retention elements identified was overlapped with the known *ecMYC* coordinates: chr8:127437980-129010086 (hg19). To analyze chromatin interactions with chromosome bookmarked regions, we used previously identified bookmarked regions that retained accessible chromatin throughout mitosis in single-cell ATAC-seq data of L02 human liver cells^38^ and filtered out regions that overlap with the known *ecMYC* coordinates as well as other *ecMYC* co-amplified regions: chr6:247500-382470, chr8:130278158-130286750, chr13:28381813-28554499, chr16:32240836-32471322, chr16:33220985-33538549. The resulting *ecMYC* retention elements and chromosome bookmarked regions were used as anchors to measure pairwise interactions via aggregated peak analysis (APA), using the .hic files in Juicer (v.1.22.01) and the “apa” function with 5-kb resolution and the following parameters: “-e -u”. Summed percentile matrices of pairwise interactions from “rankAPA.txt” were reported. Analyses for the *EGFR* ecDNA in the GBM39 cell line were performed in the same manner, using ecDNA coordinates: chr7:54830901-56117000 (hg19).

To analyze interactions between ENCODE-annotated classes of regulatory sequences, retention elements overlapping with “dELS”, “PLS”, or “pELS” annotations were categorized as distal enhancers, promoters, or proximal enhancers, respectively; those overlapping with both “pELS” and “PLS” annotations were categorized as promoters; those overlapping with both “pELS” or “dELS” annotations were categorized as proximal enhancers. To extract Hi-C read counts corresponding to interactions between different classes of elements on ecDNA and chromosomes, the Juicer Tools^83^ (v.1.22.01) dump command was used to extract read count data from the .hic files with 1-kb and 5-kb resolution using “observed NONE”. The resulting outputs were converted into GInteractions objects using the InteractionSet (version 1.14.0) package in R. To remove chromosomal regions with elevated signal due to copy-number changes (and not occurring on ecDNA), we filtered out chromosomal regions that overlap with copy-number-gain regions identified in WGS of COLO320DM using the ReadDepth (version 0.9.8.5) package. GInteractions objects containing Hi-C read counts between genomic coordinates in 1-kb resolution were overlapped with a GInteractions object containing pairwise interactions between chromosome bookmarked regions and *ecMYC* retention elements using the findOverlaps function in the InteractionSet package in R. Resulting read counts of these pairwise interactions were used to calculate read counts per kb using this formula: read counts per kb = 1000 × read counts / size of retention element bin in bp. Read counts per kb of each combination of interactions between different classes of elements were summed and divided by the total number of pairwise interactions belonging to each combination of interactions to obtain read counts per kb per interaction.

### Curation of candidate bookmarking factors

Candidate bookmarking factors were curated from three recently published studies: Raccaud et al.^40^, Yu et al.^38^, and Ginno et al.^84^ Candidate bookmarking factors identified in Raccaud et al. were identified in mouse cells; their orthologs were identified using the Mouse Genome Informatics database (http://www.informatics.jax.org/downloads/reports/HOM_MouseHumanSequence.rpt) and those not annotated as “Depleted” on mitotic chromosomes were included. Candidate bookmarking factors identified in Yu et al. were identified based on single-cell ATAC-seq analysis of mitotic chromosomes. Finally, candidate bookmarking factors identified in Ginno et al. were selected by focusing on protein factors which meet the following criterion: log2[ (C + 1) / (P + 1) ] > 0, where C denotes the mean protein enrichment values in mitotic cells from fractionated chromatin (chromatome), and P denotes the mean protein enrichment values in the proteomes of mitotic cells.

### Importance analysis of bookmarking factors

To interrogate whether retention elements contain binding sites of some bookmarking factors disproportionately more than others, we computed importance scores in R for each bookmarking factor in explaining the observed set of retention elements. First, we generated 1000 random permutations of the top 20 most enriched bookmarking factors within retention elements compared to random intervals. For each permuted list, we computed the incremental number of retention elements explained by (containing binding sites of) each bookmarking factor in the cumulative distribution. The mean of this value across all permutations represents the importance score for each bookmarking factor.

### CRISPR/Cas9 knockouts of bookmarking factors

Cas9-gRNA ribonucleoprotein (RNP) complexes were first assembled for each gRNA by mixing 30 µM gRNAs (Synthego) targeting CHD1, SMARCE1, and HEY1 as well as 2 non-targeting control gRNAs (2 separate guides per target; guide sequences are provided in **Supplementary Table 1**) separately with 20 µM SpCas9 2NLS Nuclease (Synthego) at a 6:1 molar ratio. Complexes were then incubated for 10 min at RT. Briefly, COLO320DM cells were counted, centrifuged at 300g for 5 min, and washed twice with PBS before resuspension in Neon Resuspension Buffer to a density of 4.2 × 10^5^ in 7 µl of buffer. 7 µl of cell suspension and 7 µl of RNP were mixed and electroporated per reaction according to the manufacturer’s instructions using a 10 µl Neon pipet tip under the following settings (1700 V, 20 ms, 1 pulse). Three electroporation reactions were plated for each replicate (2 per condition) into 6-well plates in 3 mL of media per well.

### Immunofluorescence staining-DNA FISH of KO mitotic cells

About 1M of cells were seeded onto 22×22cm poly-d-lysine coated coverslips two days after transfection. Next day, the cells were washed once with 1X PBS and fixed with 4% paraformaldehyde for 10 minutes at room temperature, followed by permeabilization with 1X PBS-0.25% Triton-X for 10 minutes at room temperature. Samples were blocked in 3% BSA diluted in 1X PBS for 1 hour at room temperature, followed by an overnight incubation at 4°C with primary antibodies: Aurora B Antibody (Novus Biologicals, NBP2-50039; 1:1000), CHD1 (Novus Biologicals, NBP2-14478; 1μg/mL), HEY1 (Novus Biologicals, NBP2-16818; 1:1000), SMARCE1 (Sigma-Aldrich, HPA003916; 1μg/mL). Cells were washed in 1X PBS and incubated with fluorescently conjugated secondary antibodies (F(ab’)2-Goat anti-Rabbit IgG (H+L) Cross-Adsorbed Secondary Antibody, Alexa Fluor™ 488 (Invitrogen, A-11070), Donkey anti-Mouse IgG (H+L) Highly Cross-Adsorbed Secondary Antibody, Alexa Fluor™ 647 (Invitrogen, A-31571) at 1:500 for 1 hour at room temperature. The samples were then washed in 1X PBS, and fixed with 4% paraformaldehyde at room temperature for 20 mins. A subsequent permeabilization using 1X PBS containing 0.7% Triton-X and 0.1M HCl was performed on ice for 10 mins, followed by acid denaturation for 30 minutes at room temperature using 1.9M HCl. The samples were then washed once with 1X PBS and then 2X SSC, followed by washes with an ascending ethanol concentration of 70%, 85% and 100% for 2 mins each. *MYC* FISH probes (Empire Genomics) were diluted with hybridization buffer and subjected to heat denaturation at 75°C for 3 mins, prior to applying onto the fully air-dried coverslips for overnight hybridization at 37°C. The next day, the coverslips were washed once with 0.4X SSC, then with 2X SSC-0.1% Tween 20, and counterstained with DAPI at 50ng/mL for 2 minutes at room temperature. After rinsing in ddH_2_O, the samples were air-dried and mounted onto frosted glass slides with ProLong^TM^ Diamond Antifade Mountant (Invitrogen). Samples were imaged on a Leica DMi8 widefield microscope, where z-stack images were collected and subjected to small volume computational clearing on the LAS X.

### Analysis of immunofluorescence staining-DNA FISH of KO mitotic cells

We first created a CellProfiler (version 4.2.7)^85^ analysis pipeline to quantify protein expression levels after targeted knockdown. Briefly, we split each image into four color channels (DAPI, Aurora kinase B, target protein, and ecDNA FISH), and used DAPI to segment nuclei (40-150 pixel units) with global Otsu’s thresholding (two-class thresholding). We then identified cells by starting from the nuclei as seed regions and growing outward using the protein staining signals via propagation with global Minimum Cross-Entropy Thresholding. Mean intensity of protein staining in cells was used to determine KO efficiency of target proteins compared with controls.

Next, we created a CellProfiler analysis pipeline to quantify ecDNA tethering to mitotic chromosomes after protein KO. Briefly, we identified mitotic daughter cell pairs using pairs of cells with Aurora kinase B marking the mitotic midbody as previously shown^34^. We segmented nuclei using DAPI as above and then identified cells by starting from the nuclei as seed regions and growing outward using the protein staining signals via propagation with three-class global Otsu’s thresholding (with pixels in the middle intensity class assigned to the foreground). We separately identified ecDNA foci as primary objects using adaptive Otsu’s thresholding (two-class) and intensity-based de-clumping. Masks were then created for ecDNA foci overlapping with nuclei (with at least 30% overlap) and ecDNA foci overlapping with cytoplasm (with at least 70% overlap) and defined as tethered and untethering ecDNA, respectively. The sum of pixel areas was calculated for each group of ecDNA foci and used to calculate tethered ecDNA fractions.

### Evolutionary modeling of ecDNAs

To simulate the effect of retention and selection on ecDNA copy-number in growing cell populations, we implemented a new forward-time simulation in Cassiopeia^86^ (https://github.com/yoseflab/cassiopeia). The simulation framework builds off of the forward-time evolutionary modelling previously described^6^. Specifically, each simulation tracked a single ecDNA’s copy-number trajectory and was initially parameterized by (i) initial ecDNA copy-number (denoted as *k_init_*); (ii) selection coefficients for cells carrying no ecDNA (*s_0_*) or at least one copy of ecDNA (*s_1_*); (iii) a base birth rate (*λ*_*base*_ = 0.5); (iv) a death rate (*µ* = 0.33); and (v) a retention rate (*v* ∈ [0, 1]) that controls the efficiency of passing ecDNA on from generation to generation.

Starting with the parent cell, a birth rate is defined based on the selection coefficient acting on the cell (*s = s_0_* or *s_1_*, depending on its ecDNA content) as *λ*_1_ = *λ_base_* ∗ (1 + *s*). Then, a waiting time to a cell division event is drawn from an exponential distribution: *t_b_* ∼ exp (−*λ*_1_). Simultaneously, a time to a death event is also drawn from an exponential distribution: *t_d_* ∼ exp (−*µ*). If *t_b_* < *t_d_*, a cell division event is simulated and a new edge is added to the growing phylogeny with edge length *t_b_*; otherwise, the cell dies and the lineage is stopped. We repeated this process until 25 time units were simulated and at least 1000 cells were present in the final population.

During a cell division, ecDNAs are split amongst daughter cells according to the retention rate, *v*, and the ecDNA copy numbers of the parent cell. Following observations of ecDNA inheritance previously reported^5^, ecDNA is divided into daughter cells according to a random Binomial process, after considering the number of copies of ecDNA that are retained during mitosis. Specifically, with *n_i_* being the number of ecDNA copies in daughter cell *i* and *N* being the number of copies in the parental cell:

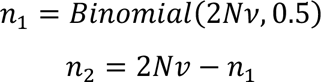

Where *Binomial* is the binomial probability distribution.

In our experiments, we simulated populations over 25 simulated time units of at least 1000 cells across ecDNA selection coefficients *s*_1_ ∈ [0, 0.8] (where *s_1_=0* indicates no selective advantage for ecDNA-carrying cells) and ecDNA retention rates *v* ∈ {0.5, 0.6, 0.7, 0.8, 0.9, 0.95, 0.97, 0.98, 0.99, 1.0} . Selection on cells carrying no ecDNA was kept at *s_0_=0*. We simulated 10 replicates per parameter combination and assessed the mean copy-number and frequency of ecDNA+ cells for each time step.

### Analysis of ecDNA sequences in patient tumors

Focal amplification calls predicted by AmpliconArchitect^87^ from tumor samples in The Cancer Genome Atlas (TCGA) and Pan-cancer Analysis of Whole Genomes (PCAWG) cohorts were downloaded from AmpliconRepository (https://ampliconrepository.org)^88^. A dataset was constructed for ecDNA, breakage-fusion-bridge (BFB), and linear amplicons containing the following information for every amplified genomic interval within each amplicon: the corresponding sample, amplicon number (within that sample), amplicon ID (assigned in AmpliconRepository), amplicon classification (ecDNA, BFB, or linear), chromosome, start and end coordinates, width, number of overlapping retention elements, and overlapping oncogenes.

Local retention element density was also computed in R for each amplified interval by dividing the number of retention elements found within 2.5 megabases of the midpoint of the interval by the local window width (5 megabases). Local retention element density was calculated for each amplicon as an average of the intervals’ local densities, weighted by interval width.

To analyze co-amplification of retention element-negative intervals with retention element-positive intervals, all amplified intervals lacking retention elements were first identified. If the amplicon corresponding to a given interval contains other intervals with retention elements, then the amplicon was considered co-amplified; each amplicon was only counted once, regardless of the number of co-amplified retention element-negative intervals. The percentage of amplicons bearing a co-amplification event was computed for each amplicon class; *p-*values were calculated between classes using a one-sided test of equal proportions.

Predicted ecDNA amplicon intervals containing *EGFR* and *CDK4*, the two most frequently amplified oncogenes within AmpliconRepository samples, were analyzed for co-amplification of oncogenes with retention elements. For each oncogene-containing ecDNA interval, 100 random oncogene-containing intervals of the same width were simulated by varying the starting point of the amplified region. For each retention element located within 500 kb of the midpoint of the oncogene’s genomic coordinates, the frequency of inclusion of that retention element within observed oncogene-containing ecDNA intervals was compared with the expected frequency based on the random intervals. Enrichment was computed as a fold-change of the observed frequency compared to the expected frequency. *P-*values comparing the distributions were calculated in R using a two-sided Fisher’s Exact Test and adjusted for multiple comparisons by the Benjamini-Hochberg method.

### DNA methylation analysis in nanopore sequencing data

Nanopore sequencing data of GBM39 was published with Zhu et al.^89^ and deposited in NCBI Sequence Read Archive (SRA) under BioProject accession PRJNA1110283. Bases were called from fast5 files using guppy (Oxford Nanopore Technologies, version 5.0.16) within Megalodon (version 2.3.3) and DNA methylation status was determined using Rerio basecalling models with the configuration file “res_dna_r941_prom_modbases_5mC_v001.cfg” and the following parameters: “--outputs basecalls mappings mod_mappings mods per_read_mods --mod-motif m CG 0 - -write-mods-text --mod-output-formats bedmethyl wiggle --mod-map-emulate-bisulfite -- mod-map-base-conv C T --mod-map-base-conv Z C”. In downstream analyses, methylation status was computed over 1 kb intervals for retention elements and other matched-size intervals within the *EGFR* ecDNA.

### CRISPRoff

CRISPRoff experiments were performed as described previously with modifications^52^. Briefly, we first cloned a plasmid (cargo plasmid) that simultaneously expresses 5 guides targeting the five unmethylated retention element sequences found on the *EGFR* ecDNA of the GBM39 cell line under U6 promoters in an array format using the previously described CARGO approach^90^ (guide sequences are provided in **Supplementary Table 1**). We also cloned a second plasmid (NTC plasmid) containing only a single *LacZ-*targeting guide, with expression also driven by a U6 promoter, as a non-targeting control. The cargo plasmid or the NTC plasmid was co-transfected with the CRISPRoff-v2.1 plasmid (Addgene, 167981) into 1.5 × 10^7^ GBM39 cells using the Neon Transfection System in accordance with the manufacturer’s protocols. Briefly, cells were dissociated to a single-cell suspension with 0.5x TrypLE, counted, centrifuged at 300g for 5 min, and washed twice with PBS before resuspension in Neon Resuspension Buffer to a density of 4.2 × 10^6^ in 70 µl of buffer; 14 µg CRISPRoff-v2.1 and 7 µg cargo or NTC plasmids were also mixed with Neon Resuspension Buffer to a total volume of 70 µl. 70 µl of cell suspension and 70 µl of plasmids were mixed and electroporated according to the manufacturer’s instructions using a 100 µl Neon pipet tip under the following settings (1250 V, 25 ms, 2 pulses). 5 electroporation reactions were pooled per replicate of each condition and cultured in T75 flasks. Cells were further cultured for two days and double positive cells (mCherry from the cargo plasmid and BFP from CRISPRoff-v2.1, or eGFP from the NTC plasmid and BFP from CRISPRoff-v2.1) were sorted using BD Aria II. The sorted cells were immediately plated on laminin-coated coverslips in a 24-well plate at a density of 1 × 10^5^ in 450 µl of media in preparation for imaging (see the “CRISPRoff imaging” section). The remaining sorted cells were cultured for an additional 3 days and harvested for genomic DNA extraction using the DNeasy Blood & Tissue Kit (Qiagen, 69504). ecDNA abundances were quantified by whole genome sequencing (WGS, see the “WGS” section).

### Imaging validation of CRISPRoff

Two days after sorting, a total of 100,000 cells were seeded onto laminin (10 µg/mL)-coated 12 mm circular coverslips for each transfection condition. Cells were allowed to recover for another 24 hours. Cells were washed once with PBS and fixed with 4% paraformaldehyde at room temperature for 10 minutes, followed by permeabilization with 1X PBS containing 0.5% Triton-X for another 10 minutes at room temperature. To further enhance fixation and permeabilization, three additional washes with Carnoy’s fixative (3:1 methanol: glacial acetic acid) were performed. The samples were then rinsed briefly with 2x SSC buffer and subjected to dehydration with ascending ethanol concentrations of 70%, 85%, and 100%. The coverslips were completely air-dried, before the application of FISH probe mixture (Empire Genomics) which was made up from 0.25 µL *EGFR* FISH probe and 4 µL hybridization buffer. The samples were denatured at 75°C for 3 minutes and then hybridized overnight at 37°C in a humidified, dark chamber. Following hybridization, the coverslips were transferred into a 24-well plate and washed once with 0.4x SSC, then 2x SSC 0.1% Tween-20, and then 2x SSC, for two minutes each. DAPI (5 ng/mL) was applied to the samples for 2 minutes to counterstain nuclei. The samples were then washed with 2x SSC and ddH_2_O prior to air dry, then mounted with ProLong Diamond. The samples were imaged on a Leica DMi8 widefield microscope using a x63 oil objective lens. *z* stacks were acquired (total range = 10 µm, step size of 0.27 µm, 38 steps) and subjected to small volume computational clearing on the LAS X software. ImageJ was used to generate maximum-intensity projections for image analysis to quantify total *EGFR* FISH copy number per nucleus.

To quantify total *EGFR* FISH copy number per nucleus, deep learning-based pixel classifiers were trained on the DAPI and *EGFR* FISH channels to create a smart segmentation and confidence mask respectively using Aivia Software (Leica Microsystems). The masks were used to create a recipe to segment FISH foci and assign FISH foci to their corresponding nucleus. The following measurements were exported for quantification: Area, Circularity, Cell.ID for nuclei; Area, Cell.ID for FISH foci. Dead cells and mis-segmented cells with a measurement in nuclei with areas greater than 200 and less than 75, and circularities less than 0.7, were excluded from the analysis. Number of cells with untethered FISH foci (i.e. FISH foci that are not within the nuclei boundaries in viable cells) were counted manually from each transfection condition.

### WGS

WGS libraries were prepared by DNA tagmentation as previously described^6^. We first transposed genomic DNA from sorted CRISPRoff cells with Tn5 transposase produced as previously described^63^, in a 50-µl reaction with TD buffer^64^, 10 ng DNA and 1 µl transposase. The reaction was performed at 50°C for 5 minutes, and transposed DNA was purified using MinElute PCR Purification Kit (Qiagen, 28006). Libraries were generated by 7 rounds of PCR amplification using NEBNext High-Fidelity 2× PCR Master Mix (NEB, M0541L) with primers bearing i5 and i7 indices, purified using SPRIselect reagent kit (Beckman Coulter, B23317) with double-sided size selection (0.8× right, 1.2× left), quantified using the Agilent Bioanalyzer 2100, diluted to 4 nM, and sequenced on the Illumina Nextseq 550. Reads were trimmed of adapter content with Trimmomatic^65^ (version 0.39), aligned to the hg19 genome using BWA MEM^66^ (0.7.17-r1188), and PCR duplicates removed using Picard’s MarkDuplicates (version 2.25.3).

### Plasmid *in vitro* methylation

To measure the effects of CpG methylation on retention element activity on a plasmid, we performed in vitro methylation of plasmids using M.SssI (NEB, M0226M) for 4 h at 37°C. Plasmids were then extracted using phenol-chloroform and precipitated using ethanol. Purified plasmids were transfected into cells and assayed using quantitative PCR or live cell imaging as described above in the sections named “Quantitative PCR analysis of plasmid retention” and “Live-cell imaging” respectively.

## Data Availability

Sequencing data generated for this study have been deposited at the NCBI SRA under BioProject accession PRJNA1333946. Coordinates (in the hg19 reference) of origins of replication identified in the K562 cell line were previously derived from SNS-seq data and published alongside those datasets at the NCBI Gene Expression Omnibus (GEO; GSE46189). GRO-seq data of COLO320DM cells were generated previously and published at the GEO (GSM7956899, replicate 1; GSM7956900, replicate 2). Asynchronous COLO320DM cell Hi-C data were previously deposited at the GEO (GSM8523315, replicate 1; GSM8523316, replicate 2). Nanopore sequencing data of GBM39 were generated in a previous study and deposited in the NCBI Sequence Read Archive (SRA) under BioProject accession PRJNA1110283. Coordinates (in the hg19 reference) of retention elements identified in the COLO320DM, GBM39, and K562 cell lines are publicly available at figshare.

## Code Availability

The ecDNA evolutionary modelling framework used in this study is publicly available through Cassiopeia^86^ at https://github.com/YosefLab/Cassiopeia.

## Materials & Correspondence

Correspondence and requests for materials should be addressed to Howard Y. Chang and Paul S. Mischel.

