## Supplementary Material for "Genetic elements promote retention of extrachromosomal DNA in cancer cells"

### Genetic elements retain extrachromosomal DNA in dividing cancer cells

**Table of Contents**

**Supplementary Table 1:** CRISPR guide RNA sequences.

**Supplementary Table 2:** PCR primer sequences.

#### Supplementary Tables

##### Supplementary Table 1.

| Guide RNA target sequence | Guide RNA information |
|---|---|
| AGCGATGCGACCCTCCGGGA | CRISPRoff cargo guide |
| TCCATCCGCCCGGTCACCGC | CRISPRoff cargo guide |
| GGCTCATCCGTGGTCGCCGG | CRISPRoff cargo guide |
| CCGCTCAACAAGTTCCCCGC | CRISPRoff cargo guide |
| CGCCATCTTGCTCGGCGCCT | CRISPRoff cargo guide |
| CCACUUGCCGUGAUAUGAAC | CHD1 KO1 |
| UUAAUUCGCCUAAGAGAACG | CHD1 KO2 |
| UCGACAGAGACAAUCUCGCA | SMARCE1 KO1 |
| UUGAUUCUCCUACCGUGACC | SMARCE1 KO2 |
| AAACCAGUCGAACUCGAAGC | HEY1 KO1 |
| AAUGUGUCCGAGGCCCGCGU | HEY1 KO2 |
| GAACGACUAGUUAGGCGUGUA | Non-targeting control 1 (NTC1) (*Gal4* targeting) |
| GTGCTGCAAGGCGATTAAGT | Non-targeting control 2 (NTC2) (*LacZ* targeting) |

##### Supplementary Table 2.

| Primer sequence | Primer information |
|---|---|
| GTCGACTCTAGAGGATCCCCTCGTCG GCAGCGTCAGATGTGTATAAGAGACAG | p5_pUC19_SmaI_20bp |
| TGAATTCGAGCTCGGTACCCGTCTCGT GGGCTCGGAGATGTGTATAAGAGACAG | p7_pUC19_SmaI_20bp |
| GTCGACTCTAGAGGATCCCC | pUC19_SmaI_5prime_fwr |
| TGAATTCGAGCTCGGTACCC | pUC19_SmaI_3prime_rev |
| TCGTCGGCAGCGTCAGATGTGTATAAGAGACAG | p5_adapter_only |
| GTCTCGTGGGCTCGGAGATGTGTATAAGAGACAG | p7_adapter_only |
| ACCATGATTACGCCAATCCAGATGCCTCTCTGGCC | pUC19_2RE forward |
| ACCTGCAGGCATGCACCTAGGCTTGAACCCCTCCA | pUC19_2RE reverse |
| GGGGTACCGAGCTCGATCCAGATGCCTCTCTGGCC | pUC19_3RE forward |
| AAACGACGGCCAGTGCCTAGGCTTGAACCCCTCCA | pUC19_3RE reverse |
| ACCATGATTACGCCAATCCAGATGCCTCTCTGGCC | pUC19_tile1 forward |
| ACCTGCAGGCATGCAGATGTGGGTGGGGCCAGATA | pUC19_tile1 reverse |
| ACCATGATTACGCCATTACAGCTCTTAAGGCGGCG | pUC19_tile2 forward |

| | |
|---|---|
| ACCTGCAGGCATGCAACACCAATCGGCACTCTGTATC | pUC19_tile2 reverse |
| ACCATGATTACGCCACCACATCCTGCTGATTGGTCC | pUC19_tile3 forward |
| ACCTGCAGGCATGCATCCACTGGGTGAAGCCAGCT | pUC19_tile3 reverse |
| ACCATGATTACGCCAGATACAGAGTGCCGATTGGTGT | pUC19_tile4 forward |
| ACCTGCAGGCATGCAGCGCTGTACTCGATTTCTCG | pUC19_tile4 reverse |
| ACCATGATTACGCCAAGCTGGCTTCACCCAGTGGA | pUC19_tile5 forward |
| ACCTGCAGGCATGCACCCTCTCTGGGCTGGCCAAG | pUC19_tile5 reverse |
| ACCATGATTACGCCAGAAATCGAGTACAGCGCCGG | pUC19_tile6 forward |
| ACCTGCAGGCATGCATGGTGAGAGGCAGAACTGGC | pUC19_tile6 reverse |
| ACCATGATTACGCCACCTTGGCCAGCCCAGAGAGG | pUC19_tile7 forward |
| ACCTGCAGGCATGCAGGCTCTGGGACTCAGCATGAGA | pUC19_tile7 reverse |
| ACCATGATTACGCCAGCCTCTTGTGCCAGTTCTGC | pUC19_tile8 forward |
| ACCTGCAGGCATGCACCTAGGCTTGAACCCCTCCA | pUC19_tile8 reverse |
| ACCATGATTACGCCAGGCATTGATTATTGACTAGT | pUC19_CMV forward |
| ACCTGCAGGCATGCAAGCTCTGCTTATATAGACCT | pUC19_CMV reverse |
| CAACAAGGTCTCCAATTGACTAGATTAGCTAGATACAGA GTGTCC | pGL4_RE1 forward |
| GCAAACGGATCCGTTTCTGGAGAAAGGGAGTAAGTAAG | pGL4_RE1 reverse |
| ATTTCCGTGTCGCCCTTATT | qPCR pUC19_F |
| ACCGCTGTTGAGATCCAGTT | qPCR pUC19_R |
| CACCGTCAAGGCTGAGAAC | qPCR GAPDH_F |
| TATACCCAAGGGAGCCACAC | qPCR GAPDH_R |
| CCGCCTCCATCCAGTCTAT | qPCR pGL4_F |
| CGAACGACGAGCGTGATAC | qPCR pGL4_R |
